# Combined loss of ATG4B, ATG4C and ATG4D impairs systemic autophagy and triggers accelerated aging in mice

**DOI:** 10.1101/2025.01.25.634746

**Authors:** Gemma G. Martínez-García, María F. Suárez, María del Mar Rodríguez-Santamaría, Víctor Celemín-Capaldi, David Roiz-Valle, Clea Bárcena, Isaac Tamargo-Gómez, Andrés Corrochano-Ruiz, Vsevolod Fedoseev, Oliva C. Fernández Cimadevilla, Pedro M. Quirós, Alejandro P. Ugalde, Pablo Mayoral, Aurora Astudillo, Christina Sveen, Nikolai Engedal, Carlos López-Otín, José M. P. Freije, Álvaro F. Fernández, Guillermo Mariño

**Affiliations:** Departamento de Biología Funcional, Facultad de Medicina, Universidad de Oviedo, Spain; Instituto Universitario de Oncología (IUOPA), Spain; Instituto de Investigación Sanitaria del Principado de Asturias (ISPA), Spain; Departamento de Bioquímica y Biología Molecular, Universidad de Oviedo, Spain; Servicio de Cardiología, Hospital San Agustín, Avilés, Asturias, Spain; Biobanco del Principado de Asturias (BBPA_ISPA_IUOPA), Registro Nacional de Biobancos PT20/161, Spain; Department of Tumor Biology, Institute for Cancer Research, Oslo University Hospital, Montebello, N-0379 Oslo, Norway; Facultad de Ciencias de la Vida y la Naturaleza, Universidad Nebrija, Madrid, Spain; Centre de Recherche des Cordeliers, Sorbonne Université, INSERM U1138, Paris, France; Centro de Investigación Biomédica en Red de Cáncer (CIBERONC), Madrid, Spain

## Abstract

Autophagy is an essential catabolic pathway that safeguards cellular and tissue homeostasis, yet the systemic consequences of its impairment in mammals remain poorly defined because complete autophagy ablation is embryonic or perinatal lethal. Here, we generate ATG4A-only mice, a model in which ATG4A is the sole remaining ATG4 protease due to combined ATG4B/C/D deletion. Through comprehensive biochemical and cellular analyses, we delineate the *in vivo* substrate specificity of ATG4A and demonstrate that it sustains only minimal ATG8 priming, uncovering a previously unrecognized functional asymmetry within the mammalian ATG4–ATG8 system. ATG4A-only mice exhibit a profound but incomplete whole-body autophagy deficiency that disrupts multiple organ systems and triggers a premature aging syndrome marked by increased DNA damage, systemic senescence, metabolic dysfunction, and dramatically shortened lifespan. Integrating these findings with comparisons to additional ATG4-deficient models, we show that organismal longevity scales with residual autophagic competence. Together, our work reveals how graded reductions in autophagy integrity influence tissue fitness and aging, establishing autophagic capacity as a key determinant of mammalian lifespan.

## INTRODUCTION

Over recent years, our understanding of autophagy has expanded substantially, establishing this conserved catabolic pathway as a central regulator of cellular homeostasis in virtually all nucleated cells. Genetic studies in cells and animal models lacking essential autophagy genes have been pivotal for defining both the core and specialized functions of the pathway. In mammals, key autophagy regulators, including *Atg3, Atg5, Atg7, Ambra1, FIP200* and *Becn1*, operate in non-redundant, central nodes of the autophagic machinery; accordingly, whole-body deletion of any of these genes causes embryonic or perinatal lethality due to complete loss of autophagy, and even tissue-specific knockouts typically result in severe dysfunction of the affected organs or cell types (1, 2). Although these models have yielded essential insights, the dramatic outcomes of full autophagy ablation limit their ability to reveal the physiological roles of autophagy under less extreme conditions. Conversely, the impact of systemically reduced, but not abolished, autophagy has been far less explored, largely due to the scarcity of appropriate genetic models. This gap is particularly relevant because autophagic activity declines progressively with age (3), and can be further compromised by environmental exposures such as pollutants or nanoparticles (4). Consistently, autophagy dysfunction has emerged as one of the defining hallmarks of aging (5, 6).

In contrast to essential autophagy genes, mammals encode four paralogues of the yeast Atg4 cysteine protease, ATG4A, ATG4B, ATG4C and ATG4D (7). All four ATG4s catalyze key reactions on the mammalian Atg8 orthologues (LC3A, LC3B, LC3C in humans, GABARAP, GABARAPL1 and GABARAPL2), collectively known as mATG8s, whose proteolytic activation is required for subsequent lipidation and autophagosome association (8). Among them, ATG4B is the predominant priming protease for all mATG8s except GABARAPL2 (9, 10), whereas ATG4D serves as the major delipidating enzyme that removes mATG8s from autophagosomal membranes once maturation is complete (11). ATG4C provides auxiliary delipidation activity, particularly in the absence of ATG4D (12), while the physiological role of ATG4A remains poorly understood. Despite these distinct biochemical preferences, partial functional redundancy within the ATG4 family allows single knockouts to develop normally (13, 14). For example, ATG4C-deficient mice display only mild, tissue-specific defects in autophagic degradation and otherwise resemble their WT counterparts (12). ATG4B-deficient mice are also viable but exhibit a significant reduction in autophagic capacity across multiple organs: autophagic flux and p62/SQSTM1 turnover are diminished under both basal and starved conditions. Although this limited autophagic competence supports a near-normal lifespan, *Atg4b^−/−^*mice fail to mount robust autophagic responses to stress and consequently show increased susceptibility to diverse experimental injuries, including bleomycin-induced pulmonary fibrosis, endotoxemia-associated lung damage, experimental colitis, optic nerve–induced cellular damage, obesity and additional metabolic disturbances (15–19). Mice deficient for ATG4D have also been recently generated. These mice develop late-onset cerebellar neurodegeneration and, although they show alterations in the status of some autophagy-relevant parameters, due to the preponderant role of ATG4D in mATG8s delipidation, do not show any alteration in autophagic flux (11).

Here, we report the generation and comprehensive characterization of mice simultaneously lacking ATG4B, ATG4C and ATG4D, which therefore express ATG4A as the sole remaining ATG4 family member (hereafter referred to as ATG4A-only). This model enables the in vivo dissection of ATG4A’s physiological substrate specificity and its contribution to autophagy in the absence of functional redundancy. Using cells and tissues from ATG4A-only mice, we delineate how ATG4B/C/D deficiency disrupts mATG8 processing and profoundly impairs multiple steps of autophagy regulation and execution. Phenotypically, ATG4A-only mice exhibit a severe but incomplete systemic autophagy defect that precipitates early-onset, multimorbid features resembling physiological and accelerated aging. These molecular alterations are accompanied by epigenetic age acceleration and by the accumulation of DNA damage, inflammation, and senescence markers, which together result in a dramatically reduced lifespan.

## MATERIALS AND METHODS

### Mouse strains

ATG4A-only mice were generated at the University of Oviedo facilities by crossing previously generated *Atg4b^-/-^*(20) *Atg4c*^-/-^(12) and *Atg4d*^-/-^(11) mice.

### Cell lines

The different mouse embryonic fibroblasts (MEF) lines that were used in this work were generated as follows: MEFs were extracted from E13 embryos. Embryos were sterilized with ethanol, washed with PBS, and triturated with razor blades. Samples were then incubated in DMEM (Gibco) overnight at 37 °C and 5% CO_2_. The next day, cultured cells were trypsinized, filtered and washed. Finally, MEFs were incubated at 37 °C and 5% CO_2_ and used for the corresponding experiments. These MEFs were maintained in Dulbecco’s modified Eagle’s medium (DMEM) supplemented with ×1 non-essential amino acids, 10 mM HEPES buffer, 100 μM 2-mercaptoethanol, ×1 sodium pyruvate and 10% fetal bovine serum. HEK293T cells were maintained in DMEM supplemented with 10% fetal bovine serum, 1% penicillin-streptomycin-l-glutamine and 1% antibiotic-antimycotic (Gibco) at 37 °C in 5% CO_2_.

### Antibodies

In this study, the following antibodies were used:

Anti-SQSTM1 (M01, clone 2C11) (Abnova Corporation, Cat# H00008878-M01); anti-LC3A (Proteintech, Cat# 12135-1-AP); anti-LC3B (Novus, Cat# NB600-1384); anti-GABARAP (MBL International, Cat# PM037); anti-GABARAPL1 (ATG8L) (Proteintech, Cat# 11010-1-AP); anti-Atg4A (Arigo, ARG54904 and Thermo Scientific, PA557632); anti-GABARAPL2 (GATE-16) (MBL International, Cat# PM038); anti- β-Actin (Sigma-Aldrich, Cat# A2228); Alexa Fluor® 488 goat anti-mouse (Invitrogen, Ref.#A-11001); Alexa Fluor® 488 goat anti-rabbit (Invitrogen, Ref.#A-11008); Alexa Fluor® 532 goat anti-mouse (Invitrogen, Ref.#A-11002); Alexa Fluor® 532 goat anti-rabbit (Invitrogen Ref.#A-11009); anti-alpha-Tubulin (Sigma-Aldrich, Cat# T5168); anti-GAPDH (Novus Cat# NB300-320); anti-ubiquitin (FK2) (Enzo Life Sciences, Cat# BML-PW8810-0500); anti-LAMP-1 (H4A3) (Santa Cruz Biotechnology, Cat# sc-20011); anti-FIP200 (D10D11) (Cell Signaling Technology, Cat# 12436); anti-ULK1 (D8H5) (Cell Signaling Technology, Cat# 8054); anti-Phospho-ULK1 (Ser555) (D1H4) (Cell Signaling Technology, Cat# 5869); anti-phospho-ULK-1 (Ser757) (D7O6U) (Cell Signaling Technology, Cat# 14202); anti-Atg101 (E1Z4W) (Cell Signaling Technology, Cat# 13492); anti-Atg13 (D4P1K) (Cell Signaling Technology, Cat# 13273); anti-ATG14/Barkor (C-Terminal) (Proteintech, Cat# 24412-1-AP); anti-Beclin-1 (Cell Signaling Technology, Cat# 3738); anti-Atg12 (Cell Signaling Technology, Cat# 2011); anti-p70 S6 Kinase (Cell Signaling Technology, Cat# 9202); anti-Phospho-p70 S6 Kinase (Thr421/Ser424) (Cell Signaling Technology, Cat# 9204); anti-4E-BP1 (Cell Signaling Technology, Cat# 9452); Anti-AMPK-alpha (Cell Signaling Technology, Cat# 2532); anti-Phospho-AMPK (Thr172) (Cell Signaling Technology, Cat# 2531); anti-NBR1 (Cell Signaling Technology, Cat# 9891); anti-NDP52 (D1E4A) (Cell Signaling Technology, Cat# 60732); anti-OPTN (Proteintech, Cat# 10837-1-AP).

### Plasmids and recombinant vectors

In this study, the following plasmids were used:

pCI-neo-myc-LC3 (deltaC22) (Addgene_45448); pMXs-IP-EGFP-ULK1 (Addgene_38193); pMXs-IP-EGFP-mAtg5 (Addgene_38196); pMRXIP-GFP-STX17TM (Addgene_45910); pMRXIP GFP-VAMP8 (Addgene_45919); pBABE-puro SV40 LT (Addgene_38193); pMXs-IP-EGFP-hAtg13 (Addgene_38191); pMXs-IP-EGFP-LC3 (Addgene_38195); pMXs-puro GFP-p62/SQSTM1(Addgene_38277). Other plasmids appearing in the paper which are not listed here were generated at our lab and are fully available upon request to the corresponding author.

### Other reagents

Other reagents and chemicals used in this study were:

Bafilomycin A1 (Enzo Life Science, Cat# BML-CM110-0100, CAS: 88899-55-2); Torin 1 (Selleckchem, Cat# S2827, CAS:1222998-36-8); 3-Methyladenine (Sigma Aldrich, Cat# M9281, CAS: 5142-23-4); 2-mercaptoethanol (Merck, Ref.# 8057401000, CAS 60-24-2); 2-Propanol (Merck, Ref.#1096341000, CAS 67-63-0); 30% polyacrylamide solution, 37.5:1 (Bio-Rad, Ref.# 1610159, CAS 0079-6-1 and 868-63-3); Ammonium Acetate (Merck, Ref.# 1011151000, CAS 631-61-8); SAR405 (Selleckchem, Cat# S7682 CAS:1523406-39-4); Ammonium persulfate (Bio-Rad, Ref.# 1610700, CAS 7727-54-0); cOmpleteTM (Roche, Ref.# 11873580001); DAPI (Sigma, Ref.# D9542, CAS 28718-90-3); DMEM (Sigma, Ref.# D5796-24X500ML); dNTPs (Invitrogen, Ref.# 10297018); 10x DPBS (Invitrogen, Ref.# 14200-083); Earle’s balanced salt solution (Invitrogen, Ref.# 24010-043); EDTA disodium salt (Affymetrix, Ref.# 156971KG, CAS 6381-92-6); EDTA tetrasodium salt dihydrate (Affimetrix, Ref.#157001KG, CAS 10378-23-1); Lipofectamine® (Invitrogen, Ref.# 18324-012); Opti-MEM (Invitrogen, Ref.# 31985-062); Paraformaldehyde solution (Merck, Ref.# 1.04002.1000, CAS 50-00-0 and 67-56-1); PEG8000 (Sigma, Ref.# P2139-1KG, CAS 25322-68-3); Pen-Strep-Glut (Invitrogen, Ref.# 10378-016); PhosSTOP (Roche, Ref.# 4906837001); Platinum Taq DNA polymerase (Invitrogen, Ref.#10966-034); Polybrene (Santa Cruz, Ref.# sc-134220, CAS 28728-55-4); Proteinase K (Roche, Ref.# 03115879001, CAS 39450-01-6); Puromycin dihydrochloride (Gibco, Ref.# a11138-03, CAS 58-58-2); RNase A (Qiagen, Ref.# 19101, CAS 9001-99-4); RNaseOUT (Invitrogen, Ref.# 10777-019); Sodium dodecyl sulfate (Merck, Ref.# 8220501000, CAS 151-21-3); Super Script® IV Reverse Transcriptase (Invitrogen, Ref.#18090050); TEMED (Affymetrix, Ref.# US76320-100GM, CAS 110-18-9); ThremoScript^TM^ RNase H (Invitrogen, Ref.#12236-022); TRizol® (Ambion, Ref.#15596018); Trypsin-EDTA (0.05%) (Gibco, Ref.# 25300-054); Tween-20 (Sigma-Aldrich, Ref.#P1379-1L, CAS 9005-64-5); Sodium pyruvate (Invitrogen, Ref.# 11360-039); MEM NEAA (Invitrogen, Ref.# 11140-035); HEPES buffer (Invitrogen, Ref.# 15630-056).

### Lentiviral transduction of MEF cells

Transfections were carried out in cells seeded onto gelatinized coverslips and using Lipofectamine, following the manufacturer’s instructions. For lentiviral infection, HEK293T cells were transfected with lentiviral vector together with packaging plasmids psPAX2 and pMD2.G using Lipofectamine. After 48 h of transfection, supernatants were filtered through 0.45 μm polyethersulfone filters to collect the viral particles and added at 1:3 dilution to previously seeded mouse fibroblasts supplemented with 0.8 µg/ml of polybrene. Selection with puromycin (2 µg/mL) was performed 2 days after infection.

### Quantitative real-time PCR

cDNA was synthesized using 1 to 5 µg of total RNA, 0.14 mM oligo (dT) (22-mer) primer, 0.2 mM concentrations each of deoxynucleoside triphosphate and SuperScript II reverse transcriptase (Invitrogen). Quantitative RT-PCR (qRT-PCR) was carried out in triplicate for each sample using 20 ng of cDNA, TaqMan Universal PCR Master Mix (Applied BioSystems) and 1 µl of the specific TaqMan custom gene expression assay for the gene of interest (Applied Biosystems). To quantify gene expression, PCR was performed at 95 °C for 10 min, followed by 40 cycles at 95 °C for 15 s, 60 °C for 30 s and 72 °C for 30 s using an ABI Prism 7700 Sequence Detection System. As an internal control for the amount of template cDNA used, gene expression was normalized to the mouse β-actin gene using the Mouse β-actin Endogenous Control (VIC/MGB Probe, Primer Limited). Relative expression was calculated as RQ=2^-ΔΔCt^.

### Protein extract preparation

Tissues were immediately frozen in liquid nitrogen after extraction and homogenized in a 20 mM Tris buffer pH 7.4, containing 150 mM NaCl, 1% Triton X-100, 10 mM EDTA and Complete® protease inhibitor cocktail (Roche Applied Science). Then, tissue extracts were centrifuged at 12.000 rpm at 4 °C and supernatants were collected. Protein concentration was quantified by the bicinchoninic acid technique (BCA protein assay kit, Pierce Biotechnology, 23225). For protein extracts derived from cultured cells, cells were washed with cold PBS and lysed in a buffer containing 1% NP-40, 20 mM HEPES (pH 7.9), 10 mM KCl, 1 mM EDTA, 10% glycerol, 1 mM orthovanadate, 1 mM phenylmethanesulfonyl fluoride (PMSF), 1 mM dithiothreitol, 10 µg/ml aprotinin, 10 µg/ ml leupeptin and 10 µg/ml pepstatin. Lysates were centrifuged at 12.000 rpm at 4 °C and supernatants were collected. Protein concentration was quantified by the bicinchoninic acid technique (BCA protein assay kit, Pierce Biotechnology, 23225).

### Immunoblotting

A total of 25 µg of protein sample was loaded on either 8% or 13% SDS-polyacrylamide gels. After electrophoresis, gels were electrotransferred onto polyvinylidene difluoride (PVDF) membranes (Millipore), and then membranes were blocked with 5% non-fat dried milk in PBT (phosphate-buffered saline with 0.05% Tween 20) and incubated overnight at 4 °C with primary antibodies diluted in 3% non-fat dried milk in PBT. After three washes with PBT, membranes were incubated with the corresponding secondary antibody at 1:10.000 dilution in 1.5% milk in PBT and were developed with Immobilon Western Chemiluminescent HRP substrate (Millipore, P36599A) by using an Odyssey® Fc Imaging System (LI-COR, Lincoln, NE, USA). Unless otherwise specified, immunoblotting against β-actin was used as sample processing control (LOAD) for the immunoblots shown in this article.

### GFP-LC3 degradation assay by flow cytometry

Primary MEFs at 80% confluency from stably expressing GFP-LC3B WT and ATG4A-only mice were incubated in Dulbecco’s modified Eagle’s medium (DMEM, Sigma-Aldrich) complemented with heat-inactivated fetal bovine serum as control condition, in amino acid-free Earle’s balanced salt solution (EBSS, Sigma-Aldrich) to induce autophagy and in EBSS with Bafilomycin A1 (50 nM, Enzo Life Science) to block the autophagic flux. After 4 h of incubation, cells were trypsinized, pelleted by centrifugation, washed with Dulbecco’s Phosphate Buffered Saline (DPBS), and pelleted again. Cell pellets were resuspended in DPBS supplemented with 5% of heat-inactivated FBS to a density of 10^5^ cells/mL, kept in ice and GFP fluorescence intensity was immediately analyzed by Fluorescence-Activated Cell Sorter (BD FACSAria II, BD Biosciences). Data analysis and graphical representation were done using the free software Flowing Software 2.5.1. The level of the GFP fluorescence intensity was normalized to the level of the control sample, set at 100%.

### Automated fluorescence microscopy

MEF cells stably expressing fluorescent markers were seeded in 96-well imaging plates (BD Falcon, Sparks, USA) 24 h before stimulation. Cells were treated with the indicated agents for 4 h. Subsequently, cells were fixed with 4% PFA and counterstained with 10 µM Hoechst 33342. Images were acquired using a BD pathway 435 automated microscope (BD Imaging Systems, San Jose, USA) equipped with a 40X objective (Olympus, Center Valley, USA). Images were analyzed for the presence of fluorescent puncta in the cytoplasm using the BD Attovision software (BD Imaging Systems). Cell surfaces were segmented and divided into cytoplasmic and nuclear regions according to standard proceedings. RB 2x2 and Marr-Hildreth algorithms were used to detect cytoplasmic GFP-LC3 positive dots.

### The lactate dehydrogenase (LDH) sequestration assay

LDH sequestration was measured as described previously (21). Briefly, WT and ATG4A-only MEFs were seeded in 6-well plates and allowed to settle for 2 days before being cultured for 3 h in complete, nutrient-rich medium (RPMI 1640/10% FBS; control) in the absence or presence of 100 nM Bafilomycin A1 (BafA1), or for 3 h in amino acid-starvation medium (EBSS) + 100 nM BafA1. Subsequently, cells were harvested (with trypsin-EDTA), pelleted at 500 x *g* at 4 °C, resuspended in 400 µL ice-cold 10% sucrose, and subjected to a single high-voltage electric pulse (2000 V and 1.2 µF in a 1 x 1 cm electrode chamber of a home-made apparatus) for selective plasma membrane disruption. After mixing with 400 µL ice-cold phosphate-buffered sucrose (100 mM sodium monophosphate, 2 mM DTT, 2 mM EDTA, and 1.75% sucrose, pH 7.5), an aliquot of 150 µL was frozen at - 80 °C for subsequent measurement of total cellular LDH levels, whilst 550 µL was mixed with 1,000 µL resuspension buffer (50 mM sodium monophosphate, 1 mM EDTA, 1 mM DTT, 5.9% sucrose, pH 7.5) supplemented with 0.5% BSA and 0.01% Tween 20, and centrifuged at 18,000 x *g* for 45 min at 4 °C. After carefully removing all supernatant (containing cytosolic LDH), the pellet (containing LDH sequestered into autophagic vacuoles) was frozen at -80 °C. Whole-cell aliquots and pellets were solubilized in 1% Triton X-405 (Sigma) in resuspension buffer, and LDH levels were quantified based on the decrease in NADH absorbance at 340 nm in a classical enzymatic reaction with 0.6 mM pyruvate and 0.36 mM NADH in 65 mM imidazole (pH 7.5), using a MaxMat PL-II multianalyzer (Erba Diagnostics, Mannheim, Germany). LDH sequestration activity was calculated as percentage of sedimentable cellular LDH in BafA1-treated cells minus the percentage of sedimentable cellular LDH in control cells cultured in complete, nutrient-rich medium in the absence of BafA1 (defined as background), divided by the incubation time with BafA1.

### Long-lived protein degradation (LLPD)

LLPD was assessed as described previously (22). Briefly, MEFs were seeded in 24-well plates in 0.5 ml RPMI 1640/10% FBS (complete medium; “CM”) containing 0.1 µCi/ml ^14^C-labeled L-valine (Vitrax, VC 308). After 3 days, cells were washed with 0.5 ml CM containing 10 mM nonradioactive L-valine (Sigma, V0513) (“CM-V”) and incubated in 0.5 ml CM-V for 18 h. Next, cells were washed with 0.5 ml EBSS (Earle’s balanced salt solution; Gibco, 24010043) amino acid starvation medium supplemented with 10 mM L-valine (“EBSS-V”), and thereafter incubated in 0.25 ml EBSS-V containing 0.1% DMSO, 10 µM SAR405, or 100 nM bafilomycin A1 (BafA1) for 3 h. Subsequently, the plates were placed on ice for 2 min, and 50 µl ice-cold PBS/2% BSA and 200 µl ice-cold 25% Trichloroacetic acid (TCA) were added to each well to precipitate cellular proteins. After overnight shaking at 4 °C, all solution from each well was transferred to Eppendorf tubes and centrifuged at 5 000 x *g* for 10 min at 4 °C. The supernatants (TCA-soluble fractions) were transferred to scintillation tubes with 5 ml Opti-Fluor scintillation liquid (PerkinElmer Life Sciences) and vortexed. The pellets and the insoluble material left in the wells (together making up the TCA-insoluble fractions) were solubilized by adding 250 µl 0.2 M KOH to each and agitating on roller (pellets) or shaker (wells) for 1 h at room temperature. The solutions were merged into scintillation tubes and mixed with 5 ml Opti-Fluor scintillation liquid by vortexing. Radioactivity was measured in a liquid scintillation counter (Packard Tri-Carb 2100TR). LLPD rate was calculated as the percentage of radioactivity in the TCA-soluble fraction relative to the total radioactivity in the TCA-soluble and -insoluble fractions, divided by the 3 h LLPD sampling incubation time. SAR405- and BafA1-sensitive LLPD (indicating autophagic-lysosomal protein degradation) was calculated by subtracting the LLPD rates measured in cells that had been incubated with EBSS-V containing 10 µM SAR405 or 100 nM BafA1 from that measured in cells that had been incubated with EBSS-V containing 0.1% DMSO.

### Immunofluorescence and histology analyses of mouse tissue sections and cells

For immunofluorescence analyses with tissue cryo-sections, sections were pretreated for 30 min in 1% H_2_O_2_/PBS, followed by 1 h in blocking solution and incubated overnight with primary antibodies. Peroxidase activity was developed with the Elite Vectastain kit (Vector Laboratories) using diaminobenzidine (Dako). Sections were coverslipped with PermaFluor Aqueous Mounting Medium (Thermo Scientific). Digital images were captured with a Nikon Eclipse 80i optical microscope using the software NIS-Elements Basic Research. For paraffin sections, slides were deparaffinized and rehydrated. Slides or wells were blocked in 10% goat serum for 10 min, incubated with primary antibodies overnight at 4 °C, washed in PBS, incubated for 40 min with secondary antibodies, thoroughly washed in PBS, and stained with DAPI for nuclear staining. For immunofluorescence analyses of MEFs, cells were grown on 96-well black clear tissue culture-treated plates, washed in PBS, and fixed in 4% paraformaldehyde in PBS at room temperature for 10 min. Primary antibody was diluted 1:100 in PBS, and incubated overnight at 4 °C. Samples were washed 3 times in PBS for 15 min each. Secondary antibody was diluted 1:300 in PBS, and incubated at room temperature for 1 h. Samples were washed 3 times in PBS for 15 min each and analyzed by fluorescence microscopy.

### Light and fluorescence microscopy analyses

IHC samples were analyzed in a Leica DM4 B Upright Microscope with a Leica DFC 7000 T camera. Images were processed with LAS X software (Leica Application Suite X software, Leica Microsystems). Fluorescence microscopy images were acquired on an Axio Observer Z1 platform with a Plan-Apochromat 40X/1.3, (working distance, 0.21 mm) equipped with an ApoTome.2 system and an Axiocam MRm camera (from Carl Zeiss, Jena, Germany). Zeiss Immersol® immersion oil was used for all microscopic analyses. ImageJ (National Institutes of Health, NIH, v. 1.53e) was the software used for image analysis and quantification.

### Proteinase K protection assay

Cells were seeded in 6-well plates and autophagy induction was done by exposure to EBSS medium for 4 h. Induction was stopped by trypsinization, and cells were washed with 1x DPBS and ruptured in homogenization buffer (20 mM HEPES at pH 7.6, 220 mM mannitol, 70 mM sucrose and 1 mM EDTA pH 8) by giving 10 strokes with a 27-gauge needle and a syringe (performed in ice). Samples were centrifuged at 4 °C for 5 min at 500 x g and the supernatants were again centrifuged at 7700 x g. Each pellet was resuspended in homogenization buffer, in enough volume to subdivide samples into four tubes: control (samples in the homogenization buffer only); with the addition of Triton X-100 until 0.5% final concentration; with the addition of proteinase K (Roche, Ref. 03115879001) up to a final concentration of 50 μg/ml; and with the addition of both, proteinase K and Triton. Samples were incubated for 45-60 min at 55 °C and immediately precipitated with 1:1 v/v trichloroacetic acid (TCA) at 10% in ice for 30 min. Samples were centrifuged at 12000 x g at 4 °C for 5 min and then washed twice with ice-cold acetone. Pellets were resuspended in 30 μl of Laemmli sample buffer (described in WB section), boiled at 95 °C for 5 min, and directly loaded and analyzed by western blotting.

### Protein aggregation assay

Cells stably-expressing RFP-p62/SQSTM1 were seeded in 96-well plates with DMEM medium. Puromycin was added to the medium at 10 µg/ml (final concentration) in 20 mM HEPES buffer (pH 6.2 – 6.8) for 24 h, to induce protein aggregates across the cytoplasm. Samples were fixed after 9 and 20 h of exposure to measure aggregate formation, the number of cells with aggregates, and cell viability. As a complementary experiment, autophagy induction was performed in 96-well plates. At 0, 9, and 20 h of exposure, the medium was changed to EBSS medium to induce autophagy. Wells with DMEM+puromycin were used as controls at the corresponding time intervals. After 2 h of autophagy induction, cells were fixed as previously described to quantify the decrease in the number of protein aggregates.

### Blood and plasma parameters

Animals were starved for 6 h before measurement to avoid any possible alteration in blood glucose concentrations as a result of food intake. Blood was extracted directly from the mandibular sinus after anesthetizing mice with isoflurane. In the case that mice were previously euthanized, blood was obtained by cardiac puncture. Determination of hematological parameters was performed with the Abacus Junior hematology analyzer (Diatron Kft). Determination of biochemical analytes was performed with the veterinary biochemical analyzer Skyla VB1+ (Skyla Corp. Hsinchu City, Taiwan). Levels of serum insulin, growth hormone, leptin and adiponectin were measured with ELISA kits following the manufacturers’ protocols (Merck Millipore, Rat/ Mouse ELISA kits, Ref. insulin EZRMI-13K; Ref. growth hormone EZRMGH-45K, Ref. leptin EZML- 82K; Ref. adiponectin EZMADP- 60K). Levels of plasma IGF-1 were measured with Quantikine® ELISA kit following its corresponding protocol (R&D systems, Ref. MG100, SMG100, PMG100). Mice blood samples were collected in the presence of EDTA. Samples were then centrifuged at 3000x g at 4 °C for 15 min. Plasma samples were collected in Eppendorf tubes and stored at -80 °C until further analysis. Samples aimed to measure adiponectin and IGF-1 needed to be diluted (1:1000 v/v and 1:500 v/v, respectively) before the analysis (buffers were provided in the kits). Samples were analyzed by duplicate in 96-well plates, where standard preparations for the calibration curves, quality controls, and blanks were also loaded. Absorbance was read at 450 and 590 nm in a multi-mode microplate reader (Synergy HT, Biotek) and the differences in the readings were recorded for further data processing. Dose-response curves were fitted to a 5-parameter logistic equation, and calculations were performed with GraphPad Prism 7 software.

### Analysis of bone structure

All tibia samples were scanned by high-resolution micro-computed tomography (SkyScan 1174, SkyScan, Kontich, Belgium). The small sample-holder device for µCT was used to fit the specimen with the long axis perpendicular to the floor of the specimen holder and the x-ray source. Images were obtained by 50 kV X-ray tube voltage and 800 µA. All specimens were scanned using a 0.5 mm aluminium filter and at 9.6 µm pixel size resolution. For each specimen, a series of 613 projection images were obtained with a rotation step of 0.3° and frame averaging 2 for a total 180° rotation. The scanning time for each sample was approximately 2 h using an exposure time of 5500 ms. Flat field correction was performed at the beginning of each scan. The images obtained during scanning were reconstructed using the software NRecon (SkyScan). The correction values of attenuation coefficient, beam-hardening, smoothing and ring-artifact reduction were the same in all samples. For morphometric analysis in 2D and 3D, the software provided by the manufacturer (CTAn) was used. Region of interest (ROI) was manually delimited in each of the samples. For the analysis of the diaphyseal cortical region, 100 slices were chosen. Global grayscale threshold levels for this area were between 88 and 250. For the trabecular region, a total of 150 slices were selected and adaptive grayscale threshold levels between 63 and 250 were used. The morphometric parameters examined were bone mineral density (BMD), ratio of bone volume/ tissue volume (BV/TV), trabecular thickness (Tb.Th), trabecular number (Tb.N) and trabecular separation (Tb.Sp) for the trabecular area and connectivity density among trabeculae. The parameters were measured according to the ASBMR histomorphometry nomenclature (23).

### *In vivo* analysis of motor functions

All *in vivo* analyses were performed with male or female mice aged 2 months. For all analyses, evaluators were blinded to genotype until analyses were completed. At least 6 mice were used for each experimental group in this type of analysis.

### Footprint analyses

Hindlimbs were dipped into red non-toxic paint, whereas forelimbs were blue painted. Mice were allowed to walk through a plexiglass tunnel where the floor was covered with a sheet of white paper (20×50 cm). The stride length for each mouse was analyzed.

### Tail suspension test

Mice were suspended by their tails and evaluated after limb positions were stabilized (normally within 60 seconds). Positions were photographically captured laterally to measure the angle of the upper forelimb against the body axis.

### Raised-beam test

Mice were acclimated to crossing a 100-cm-long wooden square beam (8-mm width) elevated 30 cm above a padded base. Mice were placed on the start platform and allowed to traverse the beam to the opposite end. The time needed to cross the entire length of the beam was measured and the number of paw slips was counted. One (1) point was awarded to each paw slip. Two (2) points were awarded to each mouse fall while the test was developed. Animals were videotaped while traversing the wooden square beam for a total of three trials. A blinded observer viewed videotapes and analyzed the data.

### Rotarod test

An accelerating rotarod LE8500 (LSI, LETICA) was used to evaluate the neuronal response of mice. 6 mice for each genotype were subjected to 6 rod trials. The rod accelerated from 4 to 40 rpm in 5 min and remained at maximum speed for the next 5 min. Animals were scored for their latency to fall (in seconds) for each trial and rested a minimum of 10 min between trials to avoid exhaustion and fatigue.

### Metabolic measurements

In vivo metabolic parameters (VO₂, VCO₂, and energy expenditure) were recorded using a Comprehensive Laboratory Animal Monitoring System (Oxymax CLAMS; Columbus Instruments). Mice were housed individually for 24 h with ad libitum access to food and water. Data were processed according to the manufacturer’s instructions. Energy expenditure, VO₂ and VCO₂ values were normalized by body weight, and time stamps were rounded to 10-min intervals using the lubridate package in R. To smooth short-term fluctuations in the time-series plots, a rolling mean with a 2-h window was applied. For statistical analysis, the area under the curve (AUC) of the normalized, un-smoothed data was calculated for each mouse over the entire 24-h period and separately for the day and night intervals. 24-hour AUC values were compared between genotypes using the Wilcoxon signed-rank test. Day and night AUC values were analyzed with a linear model including genotype and time interval (day or night) as fixed factors. Estimated marginal means were obtained to compare genotypes within each interval. To account for the non-independence of day and night measurements from the same animals, cluster-robust standard errors were computed using the sandwich variance–covariance estimator (vcovCR function, clubSandwich package).

### Quantification and statistical analysis

All data acquisition and analyses were performed by investigators blinded to the experimental group. For biochemical analyses, a minimum of three samples per genotype were used for each analysis, while in vivo analyses included at least six mice per genotype, unless otherwise specified. These sample sizes are sufficient to determine whether there is a biologically meaningful difference between different genotypes, given the known mouse-to-mouse variation in autophagy assessments in previous studies published over the past decade. As for in vitro studies, a sufficiently large number of cells/areas were analyzed to ensure the description of biologically meaningful differences, also following the methods from studies cited throughout the paper. Quantifications for fluorescence microscopy/IHC were performed by analyzing at least 50 cells per condition/mouse and the measurements were performed at least in 3 different mice per genotype/condition or in three replicas for the same condition unless otherwise specified. Moreover, results obtained in cells were reliably reproduced in at least three independent experiments. All experimental data are reported as mean ± SEM unless otherwise mentioned. The data from the analyses met the assumptions of the tests and the variance was similar between the experimental groups. Unpaired two-tailed Student’s t-test was used when comparing two experimental groups. The Prism program version 7.0 (Graph-Pad Software Inc.) was used for calculations and P values lower than 0.05 were considered significant.

### RNA-Seq

RNA extracted from mouse liver samples of WT and ATG4A-only mice was processed by the company Novogene to prepare an unstranded mRNA library (poly-A enriched) and later sequenced using the NovaSeq X Plus platform 6 Gb per sample were sequenced, generating 150 bp paired-end reads. The resulting FASTQ reads were trimmed of the Illumina library adapters using cutadapt (version 4.9) and transcript quantification was carried out using Salmon (version 1.10.3) and mouse GENCODE GRCm39 vM32 as transcriptomic reference. Quantification data was summarized to gene level and imported into R using the tximeta package (version 1.27.16), and afterwards differential expression analysis was conducted using DESeq2 (version 1.49.5). To study ATG4A-only vs. WT transcriptomic changes, samples were tested using the likelihood ratio test (LRT), comparing the full model — genotype + sex + age (previously scaled)— versus a reduced model, which only accounted for sex and age. The subsequent analysis was performed using the Benjamini-Hochberg p-adjusted value result of the LRT test, and the log2FC of ATG4A-only vs. WT expression. For the GSEA analysis, a “rank” variable was calculated as log_10_ (p-value) * sign (log2 FoldChange), and this gene ranking was used to compare ATG4A-only transcriptional profile to different signatures using the fgsea R package (version 1.35.8). Firstly, the Hallmarks subset of the Mouse Molecular Signatures database (MSigDB) (version 2023.2) was used to find alterations in well-defined biological processes. In addition, a compilation of gene sets derived from transcriptomic sequencing in several studies of physiological aging were used to detect common signatures in ATG4A-only mice (24–31). Furthermore, a collection of gene expression data from microarrays and transcriptome sequencing was used to compare expression profiles of different progeria models against ATG4A-only expression profile: DNA repair-deficient progeroid model *Ercc^-/Δ7^* (32); HGPS mouse model *Lmna^G609G/G609G^* (33), Cockayne Syndrome mouse model *Csb^m/m^ Xpa^-/-^* (34) and short-lived mouse model *RagC^S74N/+^* (35). Finally, senescence signatures described in multiple references (35–37) were also compared against ATG4A-only transcriptional profile.

### Epigenetic Clock analysis

For epigenetic analysis, genomic DNA from frozen liver of WT and ATG4A-only mice was extracted using the QIAamp DNA Micro Kit (Qiagen), to subsequently determine the concentration with the Qubit dsDNA HS Assay kit (Thermo Fisher Scientific). The bisulfite conversion of DNA and the use of the Infinium Mouse Methylation BeadChip was performed by the company Hologic Diagenode. The analysis of the data was conducted using an existing protocol (38). Thereby, the resulting IDAT files were imported in a R statistical environment, where the raw signal intensities were preprocessed with the NOOB (Normal-exponential Out-Of-Band) background intensity correction implemented in minfi (version 1.55.1) (39). Furthermore, p-values of all probes were determined, and those with a p-value above 0.01 in more than 10% of samples were removed. Beta values were obtained after normalization using the Beta MIxture Quantile (BMIQ) normalization procedure, correcting for the probe bias correction using the package wateRmelon (version 2.15.0) (40). Because the epigenetic array analysis was performed in two separate batches, potential batch effects were corrected using ComBat from the sva package (version 3.57.0) (https://doi.org/10.1093/bioinformatics/bts034). Beta values were then used to calculate M-values, which were subsequently employed for unsupervised hierarchical clustering of the top 20% most variable CpG sites, visualized using the pheatmap package (version 1.0.13). Furthermore, for epigenetic age predictions, we applied a previously published epigenetic clock to our beta values, following the descriptions provided by the authors (41). Once the ages were determined, the age deviation for each sample was calculated, defined as the difference between the epigenetic age and the chronological age (42).

### Data availability

RNA-Seq data have been deposited in the NCBI Gene Expression Omnibus under accession PRJEB101992. DNA methylation array data have been deposited in ArrayExpress under accession E-MTAB-16157. All other data supporting the findings of this study are available within the paper and its Supplementary Information files, or from the corresponding author upon reasonable request. Source data are provided with this paper.

## RESULTS

### Generation of *Atg4b^-/-^Atg4c^-/-^Atg4d^-/-^* mice (ATG4A-only)

To explore the specific in vivo contribution of ATG4A and the systemic consequences of a profound impairment in ATG4-dependent autophagy, we generated mice simultaneously deficient for *Atg4b*, *Atg4c* and *Atg4d*, hereafter referred to as ATG4A-only mice. *Atg4b^-/-^* animals carry a pT1 β-geo insertion in intron 1 that abolishes transcript expression (20), whereas *Atg4c^-/-^* and *Atg4d^-/-^* mice were generated by targeted replacement of exons 1–4 with a neo cassette (12), and by insertion of a PGK-Neo cassette into exon 1 (11), respectively (Fig. 1A). PCR analysis verified the homozygosity for all the mutations in ATG4A-only mice (Fig. 1B) and qPCR demonstrated complete absence of Atg4B/C/D transcripts together with a compensatory upregulation of Atg4A (Fig. 1C), which correlated with increased ATG4A protein levels in ATG4A-only mice tissues (Fig. 1D). While ATG4A-only mice followed a Mendelian inheritance ratio in a mixed C57BL/6N–129Sv genetic background, establishing the line on a pure C57BL/6N background proved extremely challenging. Breeding pairs showed poor parental behavior, with frequent pup cannibalization and a high rate of neonatal mortality. Moreover, the number of viable ATG4A-only offspring obtained was markedly lower than expected based on Mendelian segregation. Across the entire duration of this study (approximately eight years since the birth of the first ATG4A-only mouse in a pure C57BL/6N background) only about 25 individuals were successfully generated and characterized. Despite this, ATG4A-only neonates were indistinguishable from their WT littermates (Fig. 1E) and their neonatal survival was equivalent, indicating that ATG4A alone suffices to sustain the minimal autophagy required for embryonic and perinatal survival, in contrast with most autophagy-deficient models that die during development (1). However, despite their normal embryonic and neonatal development, ATG4A-only mice soon developed several phenotypical manifestations, such as reduced size (Fig. 1E), as detailed in the following sections.

**Figure 1.**
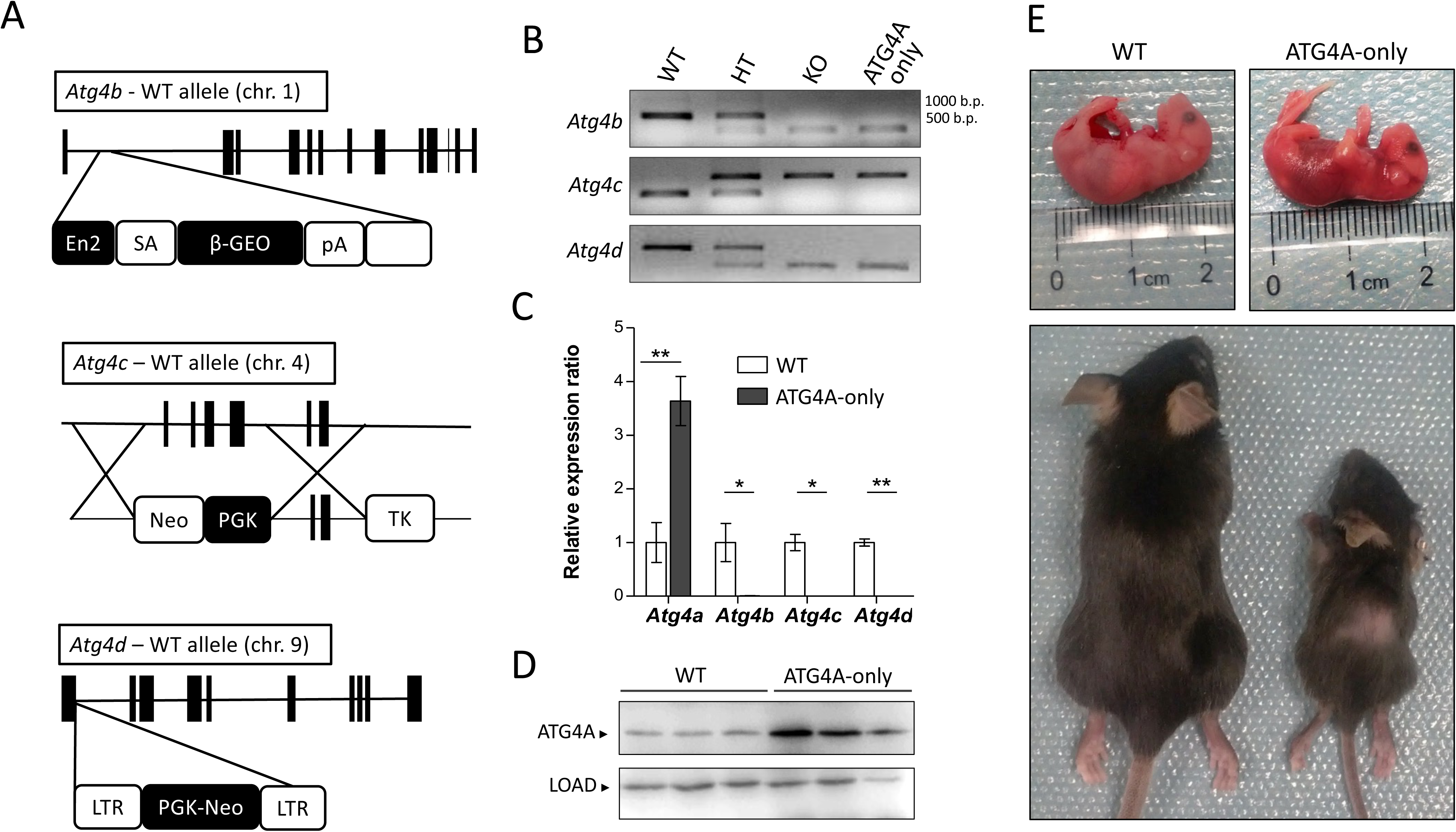
Generation of *atg4b^-/-^atg4c^-/-^atg4d^-/-^* (ATG4A-only) mice. **(A)** Schematic representations of wild-type locus for *Atg4b*, *Atg4c* and *Atg4d* genes (with coding exons represented as black boxes) showing the different gene targeting strategies to disrupt the transcription of each gene. **(B)** PCR analysis of genomic DNA from a WT control mouse (lane 1), a heterozygous control mouse (lane 2), different *Atg4* knockout control mice (lane 3, for each gene) and an ATG4A-only mouse (lane 4). **(C)** RT-qPCR analysis of kidney tissue RNA from WT and ATG4A-only mice confirming the absence of *Atg4b*, *Atg4c* and *Atg4d* expression. **(D)** Representative immunoblots of ATG4A in extracts from control and mutant mice kidneys. **(E)** Size comparison of WT and ATG4A-only neonates and 2-month-old adult mice. LOAD: β-actin. Bars represent means ± SD (N > 3 mice per genotype, at least 3 independent experiments). *, *P* < 0.05; **, P < 0.01; two-tailed unpaired Student’s t-test.

### Impact of simultaneous deficiency in ATG4B, ATG4C and ATG4D in autophagic degradation

To dissect the cellular alterations caused by combined ATG4B/C/D loss, we derived MEFs from ATG4A-only and WT embryos. Next, we performed autophagic flux experiments by subjecting WT and ATG4A-only cells to starvation including conditions in which Bafilomycin A1 (BafA1), an inhibitor of lysosomal acidification and autophagosome–lysosome fusion, was added to the culture medium. In these experimental settings WT cells showed the expected increase of lipidated mATG8 forms upon starvation, which was enhanced by BafA1 treatment (Fig. 2A). In striking contrast, ATG4A-only MEFs showed profound, substrate-specific alterations in all five mATG8s. For LC3A and LC3B, ATG4A-only MEFs accumulated a prominent high–molecular weight band corresponding to uncleaved or unlipidated species (indistinguishable under our immunoblotting conditions) and showed markedly reduced LC3A/B-II across all treatments (Fig. 2A). GABARAPL1 behaved similarly: ATG4A-only cells displayed a strong accumulation of its uncleaved form, undetectable in WT cells, together with lower levels of both unlipidated and lipidated GABARAPL1 (Fig. 2A). In contrast, GABARAP and GABARAPL2 showed a distinct pattern. ATG4A-only MEFs exhibited slightly increased GABARAP-II levels relative to WT and a robust accumulation of GABARAPL2-II under both basal and starvation conditions (Fig. 2A). Upon BafA1 treatment, LC3s and GABARAPL1-II increased only modestly, and GABARAP-II remained essentially unchanged even during starvation, indicating that autophagic flux for these substrates is almost completely blocked. Immunofluorescence analyses corroborated these findings: ATG4A-only MEFs displayed dramatically fewer LC3A, LC3B, GABARAP and GABARAPL1 puncta, whereas GABARAPL2 puncta were consistently more abundant than in WT cells under all conditions (Fig. 2B–C). To directly assess mATG8s priming efficiency, we expressed N- and C-terminally tagged LC3/GABARAP constructs in WT and ATG4A-only MEFs. ATG4A-only cells accumulated uncleaved pro-LC3s and pro-GABARAPs (Fig. 2D), whereas WT cells showed efficient processing except for a minor pool of unprocessed tagged GABARAP. As expected, GABARAPL2 processing was the least affected in ATG4A-only cells, consistent with endogenous analyses (Fig. 2A–C). The impaired autophagic flux of ATG4A-only cells translated into accumulation of autophagy substrates: p62/SQSTM1, OPTN, NDP52 and NBR1 were elevated in ATG4A-only cells (Fig. 2A), which failed to clear ubiquitin-positive aggregates or degrade GFP-p62/SQSTM1 upon starvation (Fig. 2E–G). In line with these results, FACS-based GFP-LC3B turnover assays (43) confirmed defective LC3B clearance in ATG4A-only cells (Fig. 2H). In fact, when the fusion protein GFP-LC3B is stably expressed in ATG4A-only cells, it accumulates in large p62- and ubiquitin-positive aggregates (Fig. 3A–B), consistent with inefficient LC3B priming which might tend to aggregate when present in its pro-form as previously reported for other mATG8 proteins (44). In fact, Proteinase K assays showed a higher proportion of LC3B-II associated with unclosed phagophores, indicative of defective autophagosome maturation (Fig. 3C). Functionally, ATG4A-only MEFs exhibited strongly reduced LDH sequestration and diminished degradation of long-lived proteins upon starvation in a manner that was both SAR405- (PI3KC3 inhibitor) and BafA1-sensitive (Fig. 3D). Notably, LDH sequestration was more severely impaired than long-lived protein degradation, suggesting partial compensation by non-autophagic proteolysis.

**Figure 2.**
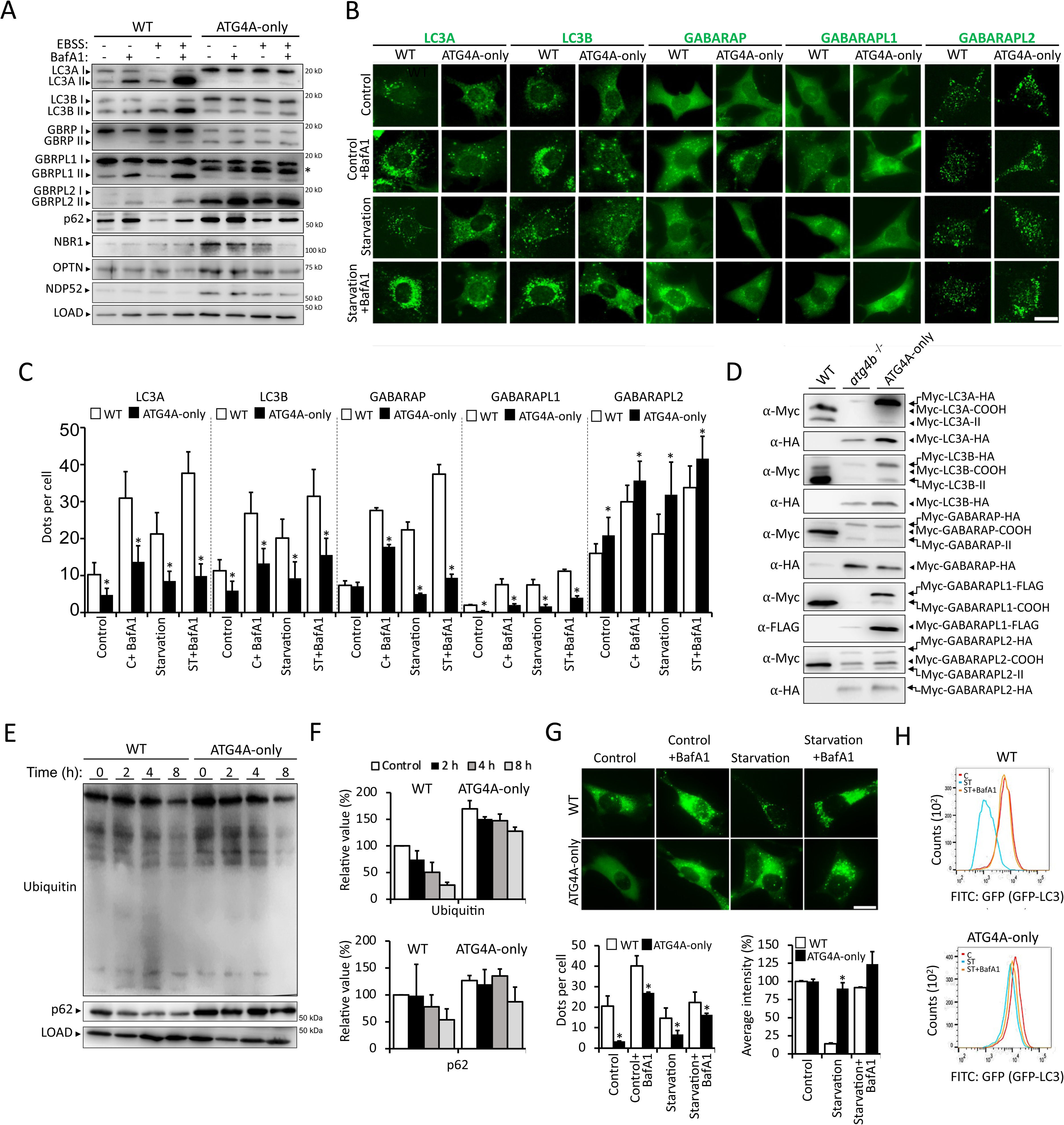
Characterization of the autophagic response in ATG4A-only cells. **(A)** Representative immunoblotting of mATG8 proteins and autophagy substrates in protein extracts from WT and ATG4A-only MEFs upon autophagy induction with EBSS medium and/or autophagy inhibition with BafA1 treatment. *, uncleaved pro-GABARAPL1 protein. **(B)** Representative images of immunofluorescence analysis of endogenous mATG8 proteins in WT and ATG4A-only MEFs. **(C)** Quantification of the data from (B). **(D)** Representative immunoblotting of protein extracts from WT, *Atg4b*-deficient and ATG4A-only MEFs that were transfected with mammalian expression vectors containing tagged cDNAs of LC3A, LC3B, GABARAP, GABARAPL1 and GABARAPL2. **(E)** Representative immunoblotting of p62/SQSTM1 and ubiquitinated proteins in protein extracts from WT and ATG4A-only MEFs after being cultured in EBSS for the indicated times. **(F)** Quantification of the data from (E). **(G)** Representative images and analysis of p62-positive structures in WT and ATG4A-only MEFs stably expressing GFP-p62. **(H)** Representative flow cytometry profiles for GFP-LC3B degradation in response to starvation in the presence/absence of BafA1 in WT and ATG4A-only MEFs. LOAD: β-actin. Bars represent means ± SEM (C, G) and ± SD (F). (N > 3 independent experiments). *, *P* < 0.05; two-tailed unpaired Student’s t-test. Scale bars: 10 µm.

**Figure 3.**
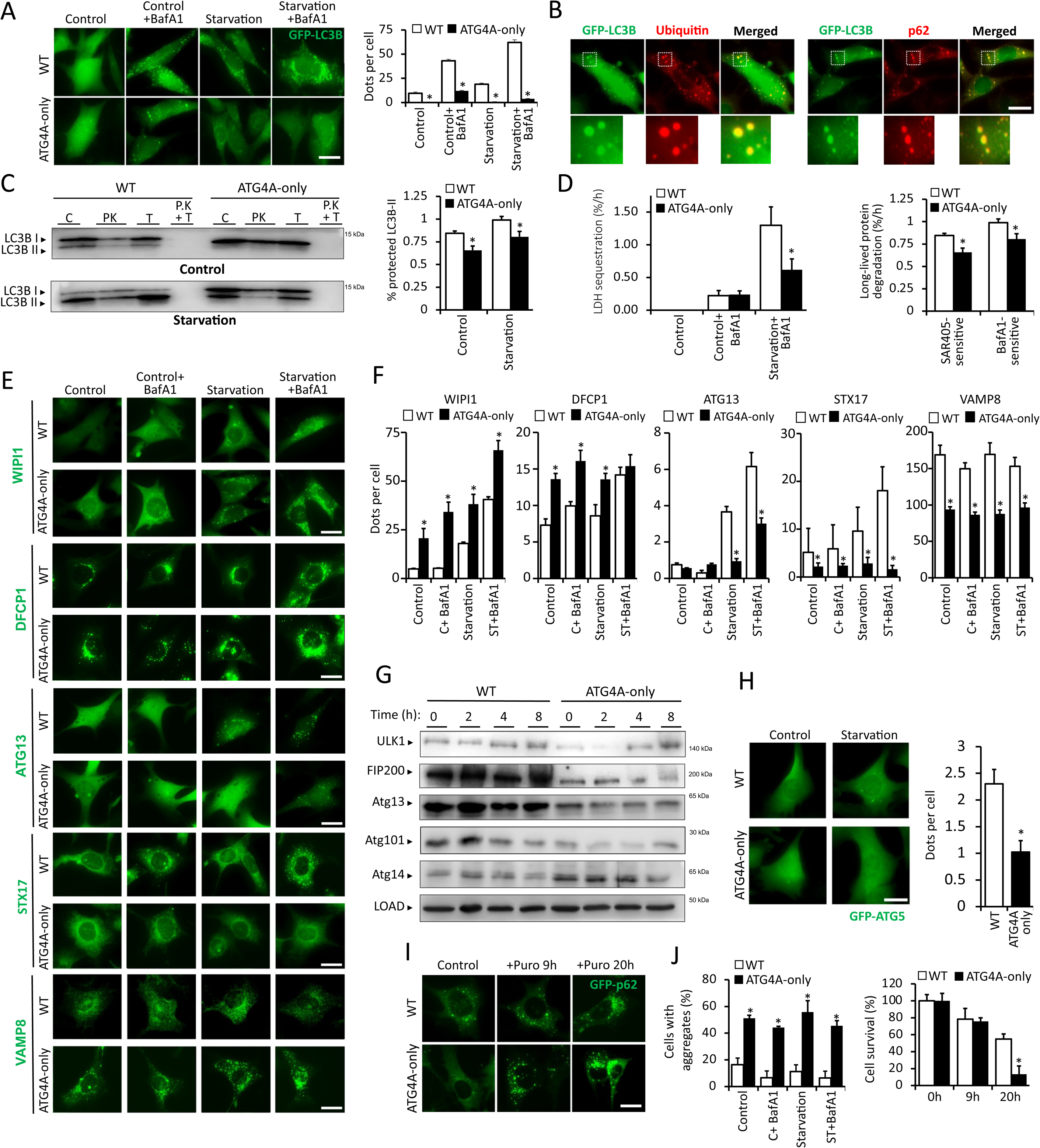
Autophagy and proteolysis regulation in ATG4A-only cells. **(A)** Representative fluorescence microscope images (left) and statistical analyses (right) of WT and ATG4A-only MEFs stably expressing GFP-LC3B upon starvation and/or BafA1 treatment. **(B)** Representative immunofluorescence images of endogenous p62/SQSTM1 and ubiquitin in WT and ATG4A-only MEFs stably expressing GFP-LC3B, showing the presence of p62- and ubiquitin-positive protein aggregates containing GFP-LC3B. **(C)** Representative immunoblotting (left) and quantitation (right) of LC3B after performing a proteinase K protection assay in WT and ATG4A-only MEFs in control or starvation conditions (C, control; PK, proteinase K; T, Triton X-100; PK+T, proteinase K + Triton X-100). **(D)** Lactate dehydrogenase (LDH) sequestration and long-lived protein degradation (SAR405- and BafA1-sensitive) assays in WT and ATG4A-only MEFs. **(E)** Representative images of WT and ATG4A-only MEFs stably expressing GFP-WIPI1, GFP-DFCP1, GFP-ATG13, GFP-STX17TM and GFP-VAMP8 in response to starvation in the presence/absence of BafA1. **(F)** Quantitation of the data from (E). **(G)** Representative immunoblotting images of components of the ULK1 complex (ULK1, FIP200, ATG13 and ATG101) and ATG14 in WT and ATG4A-only MEFs upon starvation. **(H)** Representative images of WT and ATG4A-only MEFs stably expressing GFP-ATG5 and cultured in regular or starvation medium for 4 h (left) and the quantification of the data from the autophagy-inducing condition (right). **(I)** Representative images and analysis of WT and ATG4A-only MEFs stably expressing GFP-p62/SQSTM1 after inducing the formation of protein aggregates by puromycin treatment for the indicated times. **(J)** Left, quantification of cells with aggregates after 9 h of puromycin treatment upon starvation and/or BafA1 treatment. Right, cell survival analysis of WT and ATG4A-only MEFs after the treatment with puromycin. LOAD: β-actin. Bars represent means ± SEM (N > 3 independent experiments). *, *P* < 0.05; two-tailed unpaired Student’s t-test. Scale bars: 10 µm.

Together, these findings show that ATG4A alone supports minimal priming of a subset of mATG8s—predominantly GABARAPL2—resulting in a severe block of autophagic degradation at a step prior to phagophore closure.

### Recruitment of ATG proteins in ATG4A-only cells

Autophagosome formation requires the hierarchical assembly and coordinated activity of several protein complexes that regulate autophagy initiation, phagophore expansion and autophagosome maturation (45). To investigate how the combined loss of ATG4B, ATG4C and ATG4D affects these early steps, we generated MEF lines stably expressing fluorescent reporters for key autophagy regulators and examined their basal distribution and starvation-induced recruitment in WT and ATG4A-only backgrounds. Autophagosome biogenesis relies on the orchestrated activities of the Vps34 and ULK1 complexes, which are recruited to nascent autophagosome-forming domains at the onset of autophagy (46). ATG4A-only MEFs exhibited a pronounced increase in WIPI1- and DFCP1-positive puncta (Fig. 3E–F), reflecting accumulation of PtdIns3P-enriched initiation structures generated by Vps34 (47). This phenotype resembles that previously observed in ATG5- or ATG3-deficient cells, where WIPI2 puncta accumulate due to impaired phagophore progression (45). Consistently, ATG4A-only MEFs displayed elevated levels of ATG14L, a core component of the Vps34-containing PI3KC3 complex (Fig. 3G). In contrast, recruitment of the ULK1 complex was impaired. ATG4A-only MEFs showed a clear reduction in Atg13-positive structures (Fig. 3E–F) and exhibited markedly decreased cellular levels of ATG13, ATG101 and FIP200 (Fig. 3G), all essential components of the ULK1 complex. The abundance of ATG5-positive structures—representing nascent autophagosomes (48), —was similarly reduced upon starvation (Fig. 3H), indicating defective phagophore expansion and autophagosome formation. These early defects propagated to later steps of autophagy. ATG4A-only MEFs showed a strong reduction in syntaxin 17 (STX17TM)-positive autophagosomes (Fig. 3E–F), consistent with impaired formation of fully mature autophagosomes (49). The number of VAMP8-positive vesicles, representing the lysosomal SNARE partner required for autophagosome–lysosome fusion (49). , was also markedly decreased under both basal and starvation conditions (Fig. 3E–F).Functionally, these defects rendered ATG4A-only MEFs highly vulnerable to proteotoxic stress. As previously shown (Fig. 3B), ATG4A-only cells accumulated large ubiquitin- and p62-positive aggregates upon GFP-LC3B expression. Consistently, puromycin—which induces formation of polyubiquitinated aggregates targeted for autophagic clearance via p62/SQSTM1—elicited a stronger aggregate load in ATG4A-only cells than in WT cells (Fig. 3I–J). Whereas starvation markedly reduced aggregate burden in WT MEFs after 9 h, no such reduction occurred in ATG4A-only cells (Fig. 3J). The persistent accumulation of aggregates translated into significantly impaired cell viability after 20 h of puromycin exposure (Fig. 3J).

Together, these findings demonstrate that ATG4A-only MEFs exhibit a coordinated disruption of autophagosome biogenesis: increased accumulation of WIPI1-and DFCP1-positive initiation structures, destabilization of ULK1 complex components, reduced ATG5 recruitment, and markedly impaired formation of STX17- and VAMP8-positive mature autophagosomes. This cascade of defects is consistent with the severely compromised autophagic degradation observed in these cells and explains their heightened sensitivity to proteotoxic insults requiring efficient autophagic clearance for survival.

### ATG4A-only mice show tissue-specific autophagy alterations

Given the severe autophagic defects observed in ATG4A-only MEFs, we next assessed whether these alterations extended to tissues *in vivo*. We evaluated autophagy proficiency in ATG4A-only mice by analyzing the status of mATG8 proteins and additional autophagy regulators across multiple organs. To ensure consistency, autophagy-related analyses were performed in ad libitum–fed 4-week-old WT and ATG4A-only mice with synchronized feeding–fasting cycles, as previously described (50). Immunoblotting revealed that the behavior of GABARAP, GABARAPL1 and GABARAPL2 in ATG4A-only mouse tissues largely mirrored that observed in MEFs (Fig. 4A). Specifically, GABARAP-II and GABARAPL2-II accumulated, whereas uncleaved pro-GABARAPL1 was markedly increased. In certain tissues—including skeletal muscle and lung—lipidated GABARAPL1 was also detectable (Fig. 4A). LC3 proteins showed more tissue-dependent variability: in heart, lung, kidney and liver, both LC3A and LC3B displayed the characteristic accumulation of pro-LC3/LC3-I and reduced LC3-II, similar to MEFs, whereas in brain and skeletal muscle, lipidated LC3 forms were more abundant in ATG4A-only than in WT samples (Fig. 4A). Immunofluorescence analyses confirmed these patterns. mATG8-positive puncta broadly matched the immunoblot profiles, with the exception of GABARAPL1, for which ATG4A-only tissues showed an unexpected increase in puncta across all organs examined. This may reflect accumulation of pro-GABARAPL1 aggregates or the presence of unclosed pre-autophagosomal structures retaining lipidated GABARAPL1 in the absence of ATG4D—the main mATG8 delipidating protease (11). LC3s puncta also exhibited clear tissue specificity: skeletal muscle displayed increased LC3A/LC3B puncta, whereas liver showed a marked reduction (Fig. 4B). Despite these heterogeneous mATG8 patterns, most ATG4A-only tissues exhibited substantial accumulation of the autophagy substrate p62/SQSTM1 (Fig. 4A,C). Notably, p62 accumulation in skeletal muscle and CNS regions was comparable to that in liver or heart, where LC3 lipidation was strongly reduced, indicating impaired autophagy-dependent proteostasis irrespective of LC3s-II abundance. Consistent with our findings in MEFs, ATG4A-only tissues displayed reduced levels of ULK1 complex components—ULK1, FIP200, ATG13 and ATG101—together with increased ATG14L, a marker of Vps34 complex activity (Fig. 4D). These combined defects support a model in which ATG4A-only mice experience systemic but tissue-specific disruption of autophagosome biogenesis and protein quality control.

**Figure 4:**
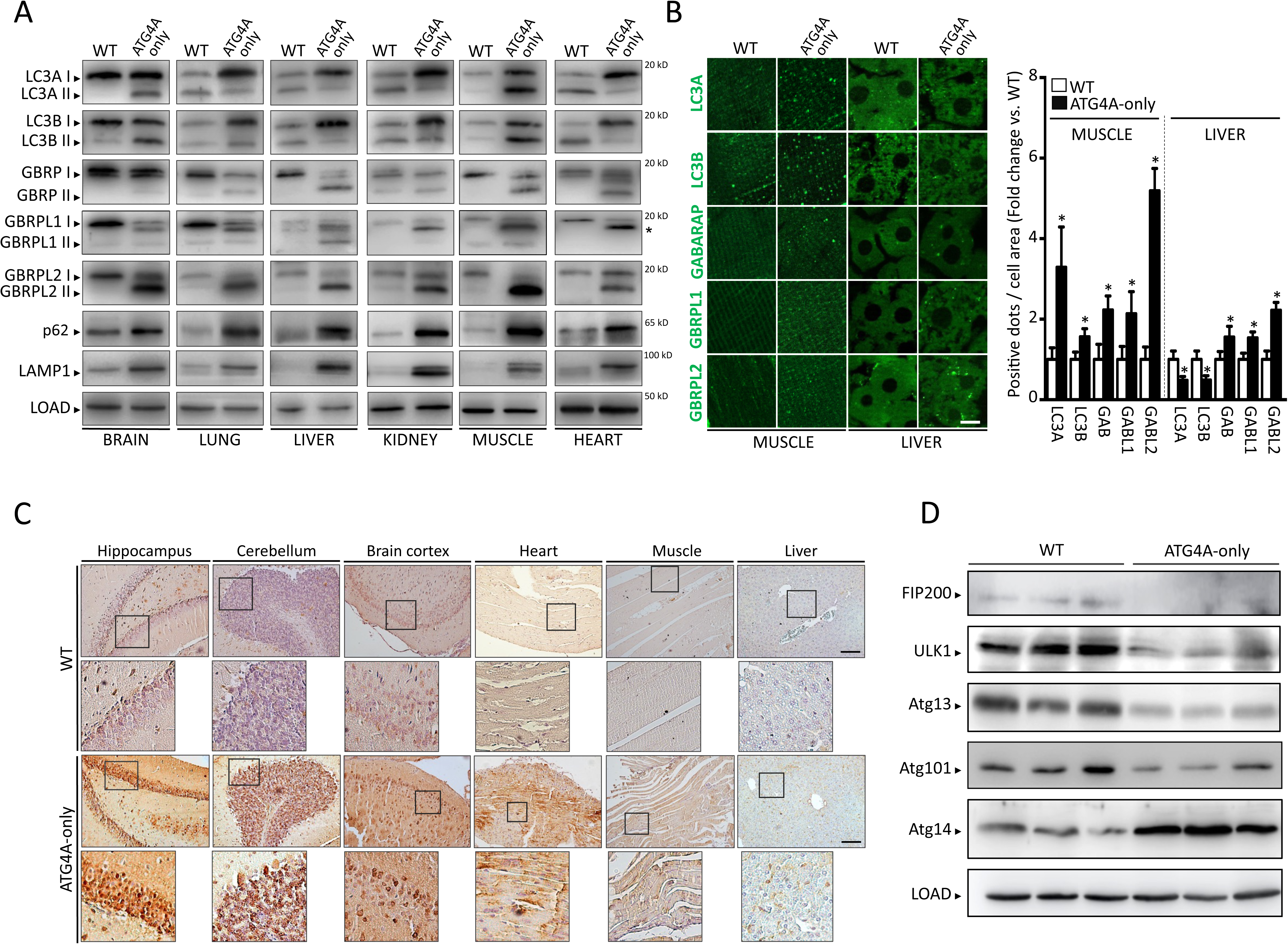
Autophagy alterations in ATG4A-only mice. **(A)** Representative immunoblots of endogenous mATG8 proteins, p62/SQSTM1 and LAMP1 in extracts from WT and ATG4A-only mice. *, uncleaved GABARAPL1 protein. **(B)** Representative immunofluorescence images (left) and quantification (right) of endogenous mATG8s in muscle and liver from WT and ATG4A-only mice. **(C)** Representative immunohistochemistry images of endogenous p62/SQSTM1 in tissues from WT and ATG4A-only mice. **(D)** Representative immunoblotting images of components of the ULK1 complex (ULK1, FIP200, ATG13 and ATG101) and ATG14 in WT and ATG4A-only mouse liver tissue extracts. LOAD: β-actin. Bars represent means ± SEM (N > 3 mice per genotype, at least 3 independent experiments). *, *P* < 0.05; two-tailed unpaired Student’s t-test. Scale bars: 10 µm (B) and 50 µm (C).

### ATG4A-only mice show reduced lifespan and features of accelerated aging

The combined loss of ATG4B, ATG4C and ATG4D profoundly disrupts the processing of most mATG8 proteins and perturbs additional components of the autophagy machinery, resulting in markedly reduced autophagic degradation in both cultured ATG4A-only cells and tissues. Consistent with these molecular defects, ATG4A-only mice developed a strong phenotype shortly after birth. Although neonatal development was normal, after weaning the mutants displayed reduced body size and weight despite normal feeding behavior (Fig. 5A, 1E) and exhibited markedly shortened survival, with a median lifespan of 133 days (range: 26–387 days) (Fig. 5B). To contextualize the severity of this phenotype, we compared ATG4A-only survival curves with those previously reported for other ATG4-deficient mice. ATG4B-deficient animals, which exhibit only a moderate reduction in autophagic activity, show limited decreases of 7.25% and 5.3% in median and maximal lifespan, respectively. Similarly, ATG4B/C- and ATG4B/D-double knockout mice display a more pronounced but still partial reduction in longevity. In striking contrast, ATG4A-only mice, harboring a profound systemic impairment in autophagy, experience the most drastic lifespan shortening among all ATG4 mutant models (Fig. 5L–N). This gradient of lifespan reduction correlates with the severity of autophagy dysfunction across genotypes, further supporting the notion that whole-body autophagic capacity is a strong determinant of organismal survival. ATG4A-only mice recapitulated many features previously described in *Atg4B*-, *Atg4C-* or *Atg4D*-deficient mice, yet in more severe form. These included pronounced balance and vestibular defects—tilted head and circling behavior—linked to impaired otoconial development (Fig. 5C and Supplementary Video 1), previously observed in *Atg4b^⁻/⁻^* mice (48). Triple mutants also developed strong ataxic manifestations, including pronounced limb-clasping (Fig. 5D), abnormal gait (Fig. 5E), reduced skeletal muscle strength (Fig. 5F), and poor performance in raised-beam and rotarod assays (Fig. 5F–G), resembling but exceeding the deficits reported in aged *Atg4D^- /-^*mice (11). Hematological analyses revealed additional alterations—thrombocytosis, reduced hematocrit, increased circulating granulocytes, and diminished mature lymphoid lineages (Fig. 5H–K)—which were more pronounced than those described for *Atg4C-*deficient mice (12), as they presented thrombocytosis, reduced haematocrit, increased circulating granulocytes and decreased mature lymphoid populations (Figs. 5H-K). Beyond these shared phenotypes, ATG4A-only mice displayed additional abnormalities not observed in single knockouts. Anatomical analyses of 3-month-old mutants revealed altered organ proportions, including increased relative size of heart, spleen, brain, pancreas and thymus, together with reduced submaxillary glands and near-complete absence of white adipose tissue (Fig. S1). Histopathology uncovered widespread structural defects (Fig. 6A): skin atrophy with hyperkeratosis, cystic dilatations and reduced subcutaneous fat; diminished brown adipose tissue; renal cortical thinning, tubular cystic dilatation and extensive fibrosis; pulmonary fibrosis; pancreatic islet reduction and focal autolysis; splenic white pulp depletion with lymphoid loss and myelopoiesis; colon wall thickening due to muscularis externa hypertrophy; and cardiac hypertrophy with fibrotic foci. Tibial µCT analysis further revealed premature bone loss with reduced bone volume, trabecular number and bone mineral density (Fig. 6B). Serum biochemical analyses indicated largely preserved homeostasis (normal sodium, phosphorus, creatinine, creatine kinase, albumin, globulins, lipase, bilirubin), but mutants exhibited increased alanine aminotransferase—consistent with liver injury—and reduced α-amylase, triglycerides, cholesterol and bile acids (Fig. 6C). Collectively, these findings highlight the essential role of autophagy in maintaining tissue integrity. Although no single catastrophic lesion explained premature lethality, the data strongly suggest multisystem functional collapse arising from compounded autophagy failure. Strikingly, many abnormalities observed in young ATG4A-only mice (including reduced muscle strength, skin atrophy, loss of subcutaneous fat, lung and kidney fibrosis, cardiac hypertrophy, lymphoid depletion and decreased bone mineral density) mirror phenotypes characteristic of aged animals and of multiple progeroid models (51). Because metabolic rewiring is a hallmark of accelerated aging (33, 34, 52, 53), we examined systemic metabolic and endocrine parameters. ATG4A-only mice showed reduced circulating leptin, consistent with their profound lipodystrophy, and markedly elevated adiponectin (Fig. 6D), a combination commonly observed in premature aging (33, 54). Dysregulation of the somatotrophic axis, a hallmark of progeroid mice (33, 34, 55), was also evident: growth hormone (GH) levels were abnormally elevated, whereas IGF-I levels were normal or reduced, consistent with hepatic GH resistance (52) (Fig. 6D). Additional aging-associated features were present, including increased hepatic glycogen deposition (34, 54) and elevated energy expenditure (linked to higher oxygen consumption and CO₂ production), metabolic traits reduced during normal aging but enhanced in progeroid animals (56–58) (Figs. 6E-F and S2).

**Figure 5:**
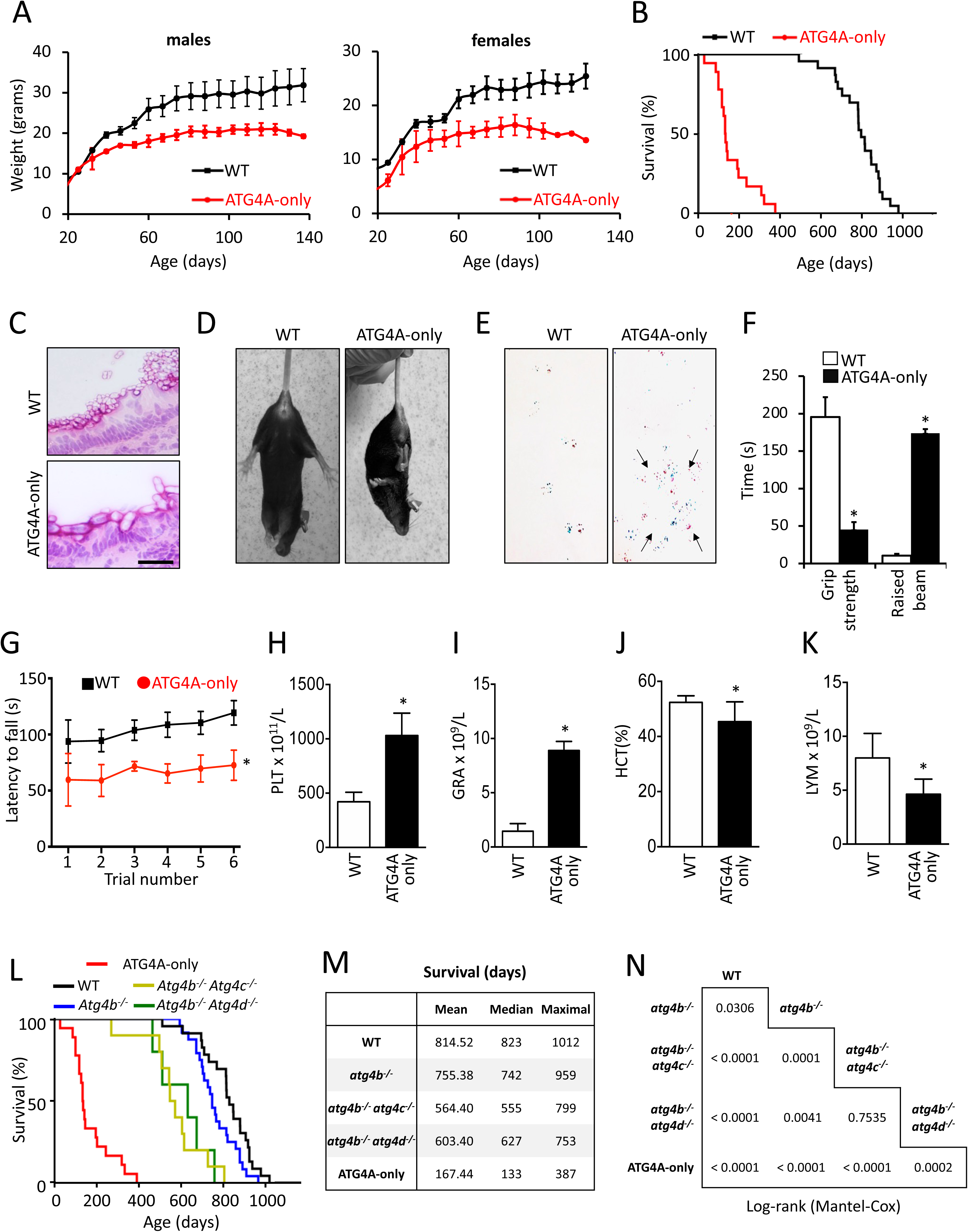
Characterization of ATG4A-only mice. **(A)** Weight comparison between WT and ATG4A-only male and female mice. **(B)** Survival comparison of WT and ATG4A-only cohorts. **(C)** Representative H&E images of the inner ear of WT and ATG4A-only mice, showing alterations in otoconial development. **(D)** Representative limb postures responding to tail suspension in WT and ATG4A-only mice. **(E)** Representative paw placement records of WT and ATG4A-only mice after footprint pattern analysis. **(F)** Performance on the raised-beam (measured by the time mice spent before reaching the end of the beam) and grip strength (measured by the time mice were able to hold themselves to a grid when inverted) tests in WT and ATG4A-only mice. **(G)** Results from Rotarod analyses of WT and ATG4A-only mice. **(H-K)** Quantification of the levels of platelets (H), granulocytes (I), hematocrit (J) and lymphocytes (K) in blood samples from WT and ATG4A-only mice. **(L)** Survival analysis of WT, *Atg4b^-/-^*, *Atg4b^- /-^ Atg4c^-/-^* and *Atg4b^-/-^ Atg4d^-/-^* and ATG4A-only mouse cohorts. **(M)** Mean, median and maximal survival values for each cohort. **(N)** Statistical analysis (Log-rank Mantel-Cox test) for all pairwise comparisons between cohorts. Bars represent means ± SD (N > 6 mice per genotype, at least 3 independent experiments). *, *P* < 0.05; two-tailed unpaired Student’s t-test. Scale bar: 50 µm.

**Figure 6:**
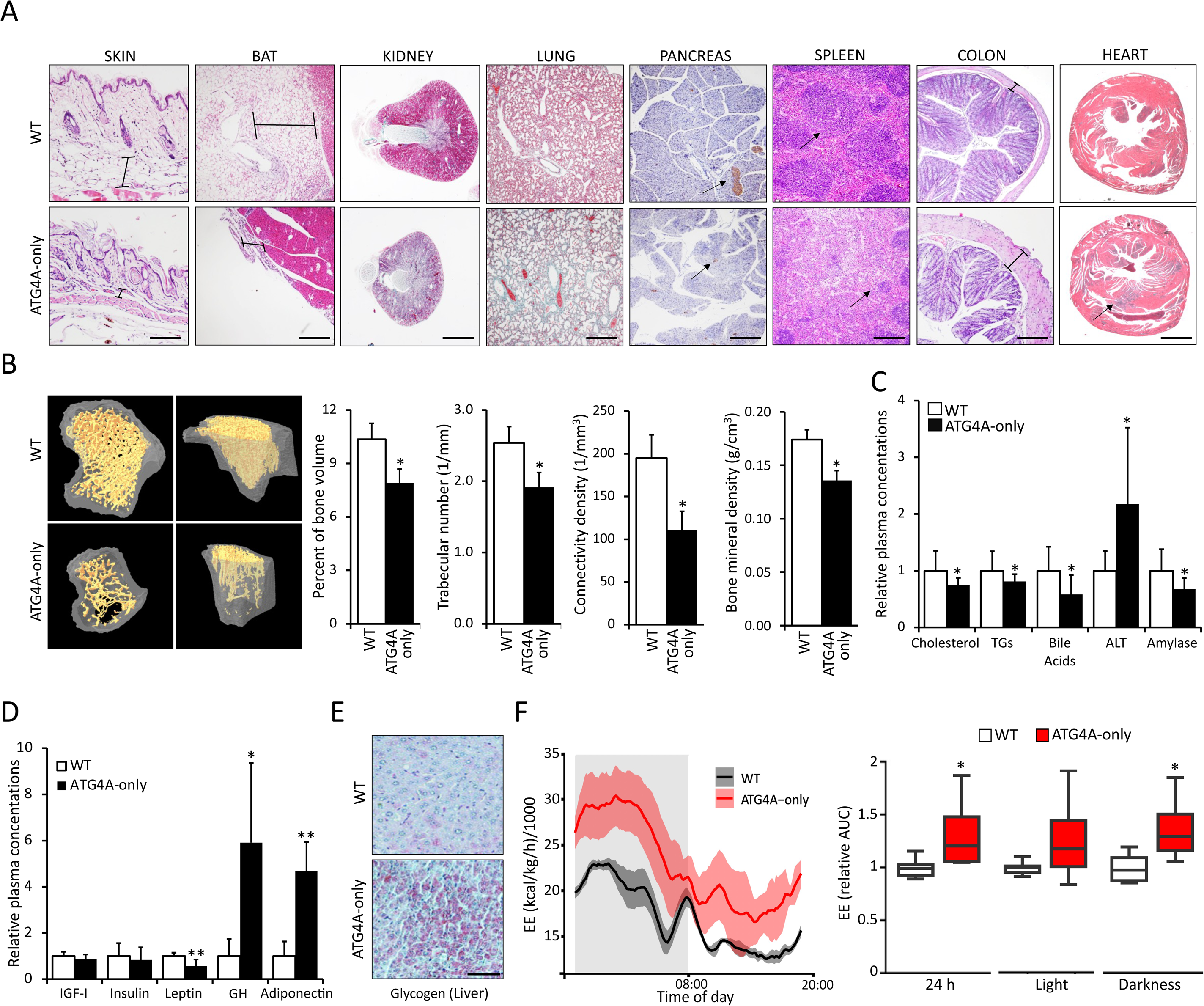
Organ-specific and biochemical alterations in ATG4A-only mice. **(A)** Representative images of H&E (skin, brown adipose tissue, spleen and colon) and Gomori trichrome (kidney, lung, heart) and anti-insulin IHC (pancreas) stainings of tissues from WT and ATG4A-only mice (arrows represent damaged/fibrotic areas). Scale bars: 200 µm (skin, BAT, lung, pancreas, spleen and colon), and 500 µm (kidney, heart). **(B)** Representative images and analysis of bones from WT and ATG4A-only mice, showing the quantification for morphometric parameters like the percentage of bone volume (ratio bone volume/tissue volume), the trabecular number, the connectivity density among trabeculae and the bone mineral density. **(C)** Quantification of major biochemical parameters (including cholesterol, triglycerides, bile acids, hepatic aminotransferases and amylase) in blood samples from WT and ATG4A-only mice. **(D)** Quantification of hormonal parameters (including IGF-1, growth hormone, insulin, leptin and adiponectin) in blood samples from WT and ATG4A-only mice. **(E)** Representative images of periodic acid-Schiff (PAS) staining in livers from WT and ATG4A-only mice showing hepatic glycogen accumulation in mutant animals. Scale bars: 50 µm. **(F)** Analysis of energy expenditure in WT and ATG4A-only mice (N = 5 and 6 respectively). Line plots represent mean ± SEM. *, *P* < 0.05; Wilcoxon signed-rank test (24 h), linear models with cluster-robust standard errors (day/night).

### Transcriptomic remodeling in ATG4A-only mice reveals inflammatory activation and metabolic suppression

Prompted by the pronounced aging-like phenotype of ATG4A-only mice, we next examined whether these systemic alterations were reflected at the transcriptional level. Bulk RNA sequencing (RNA-Seq) was performed on livers from 4-month-old WT and ATG4A-only mice. Principal-component analysis revealed clear genotype-driven segregation (Fig. S3), and differential expression analysis identified 401 significantly altered genes, 241 upregulated and 160 downregulated, in ATG4A-only livers relative to WT controls (Table S1). As expected, transcripts encoding ATG4B and ATG4C were among the most strongly reduced. Notably, 17 members of the major urinary protein (MUP) family also ranked among the most significantly downregulated genes (Fig. 7A–B, Table S1). Reduced hepatic MUP abundance and corresponding presence in urine were validated by immunoblotting and Coomassie-stained SDS–PAGE, respectively (Fig. 7C). Because MUP expression is tightly regulated by somatotroph signaling (59) and is diminished during physiological aging (60) and in progeroid animals (52), their suppression in ATG4A-only mice is consistent with systemic endocrine dysfunction. Other significantly downregulated genes included Serpinb1a, an inhibitor of IFN-driven inflammatory signaling through IKBKE degradation (61) or DIO1, an iodothyronine deiodinase essential for thyroid hormone metabolism whose activity decreases in liver injury and multiple accelerated-aging models (62, 63) (Fig. 7A–B, Table S1). Conversely, among upregulated genes, we found robust induction of pro-inflammatory mediators, including multiple CC and CXC chemokines, granzymes A and B, and the alarmin S100a9; the increased abundance of several of these proteins was confirmed by immunoblotting (Fig. 7C). This pattern is notable given that hepatocyte-specific S100a8/a9 overexpression induces Cxcl1 and drives systemic neutrophilia in mice (64). In line with this pro-inflammatory scenario, Lipocalin-2 (LCN2) was strongly upregulated at both transcript and protein levels (Fig. 7C). LCN2 is of particular interest because it can modulate LC3 lipidation through direct interaction with ATG4B (65) and is potently induced by inflammatory cytokines during liver injury (66). In line with these observations, pathway enrichment analysis showed that ATG4A-only mouse livers presented significant upregulation of several inflammatory signatures, such as interferon γ (IFNγ) and IFNα responses, tumor necrosis factor (TNFα) signaling and interleukin-6 (IL-6) signaling (Fig. 7D and S4, S6A). Moreover, we also found upregulation of allograft rejection (which involves enrichment in pro-inflammatory genes and cytokines among others), complement signaling and UV response signatures in ATG4A-only mouse livers (Fig. 7D and S4, S6A). In contrast, the most significantly suppressed pathways comprised fatty acid synthesis, peroxisomal metabolism and bile acid metabolism (Fig. 7D; Fig. S5, S6B), consistent with the lipodystrophy and reduced systemic bile acids observed in ATG4A-only mice. Taken together, the transcriptional profile of ATG4A-only livers reveals a pronounced shift toward inflammatory activation coupled with broad suppression of metabolic programs. When integrated with the observed increase in circulating granulocytes, these findings point to enhanced *inflammaging*—the chronic, low-grade, myeloid-driven inflammatory state characteristic of aging. These results reinforce the conclusion that severe systemic autophagy impairment in ATG4A-only mice is sufficient to drive widespread molecular features of aging-associated pathologies at a young age.

**Figure 7.**
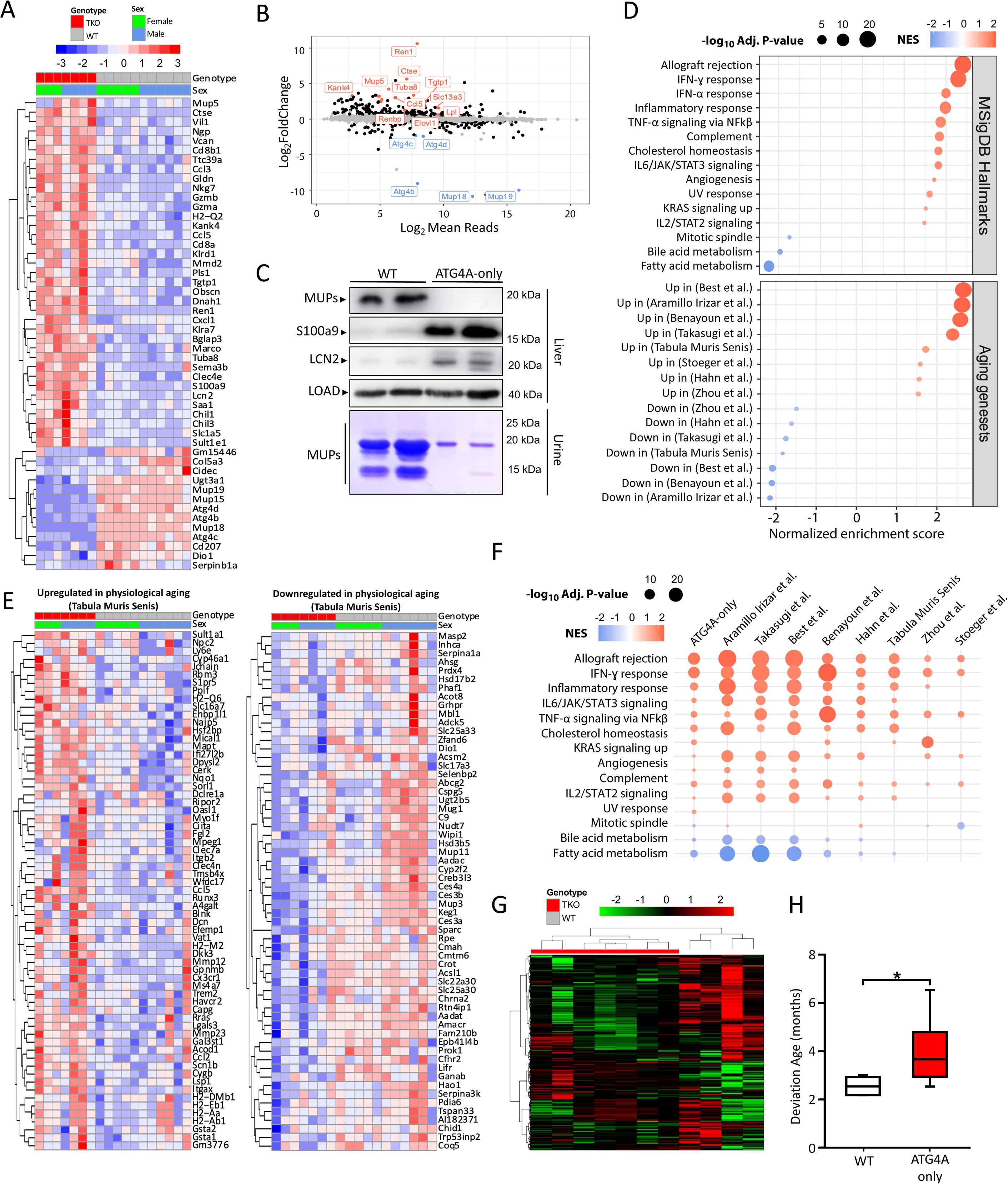
Transcriptomic analysis of ATG4A-only mice. **A)** Heatmap displaying gene expression of the top differentially expressed genes in liver ATG4A-only compared to WT mice. Significant genes were filtered to capture those with an absolute log_2_FC value greater than 2, and the top 50 genes with the lower adjusted p-value were selected for representation. **B)** MA plot showing the differential expression analysis of ATG4A-only versus WT liver mouse transcriptomes. The names of the top 20 genes with the lower adjusted p-value are indicated in the plot along with the direction of their change. **C)** Western blot analysis of MUPs, LCN2 and S100a9 protein levels in WT and ATG4A-only liver. LOAD: GAPDH. Bottom panel shows Coomassie-stained SDS–PAGE of 1 µL urine. **D)** Gene expression signatures significantly altered in ATG4A-only liver transcriptome. GSEA was performed using gene sets from MSigDB Hallmarks (up) and several transcriptomic studies of physiological aging (down). Circle size correlates with the log_10_ adjusted p-value of the analysis, whereas the color indicates the normalized enrichment score (NES) of each collection of genes, being red for upregulated pathways, and blue for downregulated pathways. **E)** Heatmap of genes in the leading edge of GSEA analysis against transcriptional profiles of physiological aging obtained from the Tabula Muris Senis consortium. The left panel shows the genes enriched in the “Upregulated genes in aging” signature, whilst the right panel shows the genes enriched in the “Downregulated genes in downregulated” collection. **F)** Summary table of GSEA results against MSigDB hallmarks between ATG4A-only mice and several transcriptomic studies of physiological aging. Circle size correlate with the log_10_ adjusted p-value of the analysis, whereas the color indicates the normalized enrichment score (NES) of each collection of genes, being red for upregulated pathways, and blue for downregulated pathways. **G)** Heatmap of the top 20 percent most variable CpG sites using unsupervised hierarchical clustering, colors ranging from green for low methylation to red for high methylation. **H)** Box plot showing the Deviation Age (DevAge) of ATG4A-only and WT mice. DevAge can be defined as the difference between epigenetic and chronological age. *, *P* < 0.05; one-tailed unpaired Student’s t-test.

### ATG4A-only mice recapitulate transcriptional and epigenetic signatures of physiological and premature aging

The pronounced inflammatory activation and metabolic suppression observed in ATG4A-only mice prompted us to investigate whether these alterations reflect molecular programs typically associated with physiological aging. Therefore, we first compared the ATG4A-only liver transcriptome with datasets from the Tabula Muris Senis consortium (31), which profiles gene expression changes during physiological aging in wild-type mice. This analysis revealed a strong positive correlation between ATG4A-only and aged WT livers, confirmed by hierarchical clustering of leading-edge genes showing nearly identical directionality of transcriptional changes (Fig. 7E). Extending this comparison to multiple independent studies of physiological aging (24–30), further revealed a strong concordance between age-associated DEGs in these datasets and those detected in ATG4A-only livers (Fig. 7D). In line with these findings, most of the pathways significantly altered in ATG4A-only mice (13 out of 15) showed concordant regulation in aged wild-type animals, underscoring a broad molecular convergence between autophagy deficiency and physiological aging (Fig 7F). Pathways commonly upregulated included TNFα, interferon-α/γ and IL6–JAK–STAT3 signaling, whereas mitochondrial, metabolic and DNA repair processes were coordinately downregulated, mirroring hallmark features of natural aging. Taken together, these extensive transcriptional similarities highlight a profound molecular correspondence between ATG4A-only and aged wild-type mice. Intriguingly, this convergence was further substantiated at the epigenetic level. In fact, unsupervised hierarchical clustering of DNA methylation profiles clearly separated ATG4A-only and WT samples, indicating a distinct epigenetic landscape associated with autophagy deficiency (Fig. 7G). In line with these observations, epigenetic clock analysis revealed that ATG4A-only mice exhibit a significantly higher biological age than WT controls, consistent with an epigenetic signature of premature aging (Fig. 7H). To determine whether this molecular aging phenotype paralleled that observed in established models of accelerated aging, we compared the ATG4A-only DEGs with datasets from different progeroid mice, including Cockayne syndrome (34), Hutchinson-Gilford progeria (33), XPF progeria (55) or *RagC^S74N/+^* mice, recently described as a new model of accelerated aging (35), yielded a striking enrichment of upregulated and downregulated genes (Fig 8A, B, S7A). Although the molecular etiologies of progeroid disorders are diverse, most involve defects in nuclear envelope stability or DNA repair, leading to hyperactivation of DNA damage responses (DDR) (67) typically mediated through p53- and/or p16-dependent pathways (68). Interestingly, when we analyzed markers of DNA damage, such as γH2AX, and the status of the main DDR effectors, such as p16 or p53, we observed a clear increase in the levels of these proteins and their phosphorylated forms in a variety of ATG4A-only mouse tissues, as compared with age-matched WT littermates (Fig. 8C). Consistently, immunohistochemistry analyses revealed nuclear accumulation of p21 in ATG4A-only mouse livers (Fig. 8D). These results show a hyperactivation of DDR in ATG4A-only mice, which likely drives the observed upregulation in the expression of inflammatory genes and increased senescence. In line with these results, when we compared the differentially expressed genes (DEGs) from our RNA-Seq experiments with datasets from mice models of senescence inflammatory responses (SIR) (36) or senescence-associated secretory phenotype (SASP) (37), we found a significant enrichment of upregulated DEGs from ATG4A-only mouse livers among those overexpressed in those datasets (Fig. 8E, S7B), reinforcing the idea of a pro-inflammatory, pro-senescent state in ATG4A-only mice.

**Figure 8:**
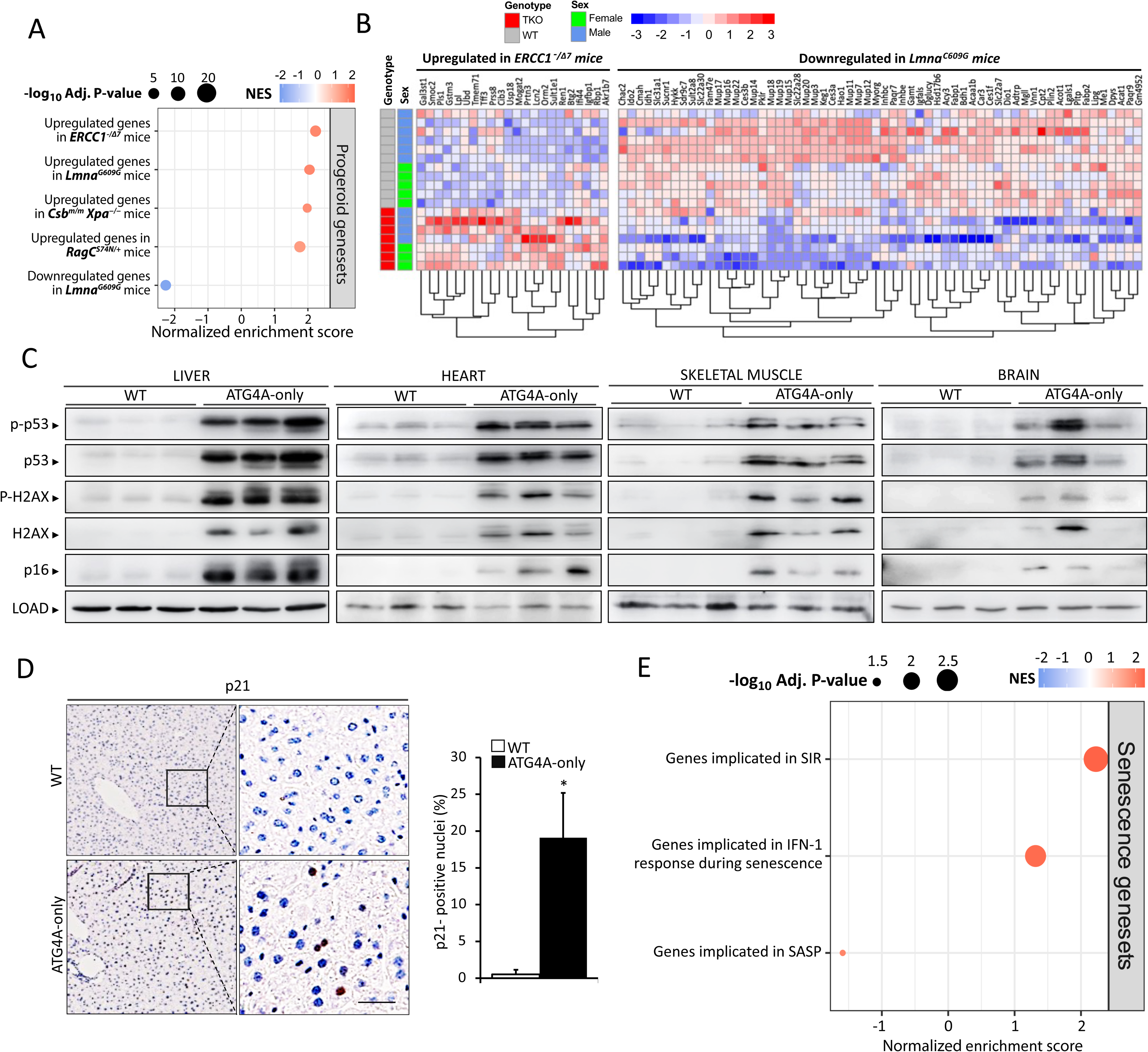
ATG4B/C/D deficiency leads to accelerated aging. **(A)** Gene expression signatures associated with progeroid models significantly altered in the ATG4A-only liver transcriptome, determined by GSEA. Circle size corresponds to the log_10_ adjusted p-value of each analysis, while circle color represents the normalized enrichment score (NES): red indicates upregulated pathways and blue indicates downregulated pathways. **(B)** Heatmap of genes in the leading edge of GSEA analysis against previously published progeroid transcriptional profiles. The left panel shows the genes enriched in the “Upregulated genes in *Ercc^-/Δ7^* mice” signature, whilst the right panel shows the genes enriched in the “Downregulated genes in *Lmna^G609G/G609G^*mice” signature. **(C)** Representative immunoblots of markers of DNA damage and DNA damage response (DDR) in tissues from WT and ATG4A-only mice. **(D)** p21 immunohistochemistry (left) and its quantification (right) in livers from WT and ATG4A-only mice. **(E)** Senescence-related gene expression signatures significantly altered in ATG4A-only liver transcriptome. SASP and IFN-I genesets were obtained from (37) and SIR geneset from (36). Circle size and color correlate with the log_10_ adjusted p-value of the analysis and NES, respectively.

Collectively, all these results show that the reduction of autophagic competency caused by the simultaneous loss of ATG4B, ATG4C and ATG4D proteases in mice leads to a progressive loss of organismal fitness, premature multimorbidity and drastically shortened lifespan, with molecular, histological, metabolic and transcriptomic alterations which are common to aged individuals and are characteristic of murine models of human premature aging syndromes.

## DISCUSSION

In this work, we provide a comprehensive dissection of ATG4 protease functions in vivo through the generation of ATG4A-only mice, a unique model in which ATG4A is the sole remaining member of the ATG4 family. This system circumvents the long-standing challenge posed by the functional redundancy of ATG4 proteases and has allowed us to define with unprecedented precision the substrate specificity of ATG4A, as well as the physiological consequences of severely reduced systemic autophagy. Our biochemical and cellular analyses reveal that ATG4A displays highly restricted priming activity *in vivo*, efficiently processing only pro-GABARAPL2. These findings contrast with earlier studies performed in overexpression systems, which suggested broader substrate specificity (69–71). Instead, our data support a scenario in which physiological ATG4A activity is narrow and insufficient to sustain normal priming of LC3A, LC3B, GABARAP and GABARAPL1 in the absence of ATG4B, ATG4C and ATG4D. Despite this deficiency, ATG4A-only cells and tissues show the presence of their lipidated forms, with an increased number of subcellular puncta positive for GABARAP proteins in most mouse tissues. We attribute this effect to the combined loss of ATG4D, the primary mATG8 delipidase (11), and ATG4C, which contributes to delipidation when ATG4D is absent (12). This delipidation impairment likely causes the small fraction of GABARAP and GABARAPL1 that is primed by ATG4A (and subsequently lipidated) to remain conjugated to membranes, as previously proposed (72). This is supported by the fact that GABARAP-II and GABARAPL1-II levels in ATG4A-only cells are higher than those observed in *Atg4B* KO cells, but similar to those observed in *Atg4B/D* double mutants (Fig. S8). Notably, LC3 lipidation showed a pronounced tissue-dependent pattern: LC3-II levels were reduced in most organs but unexpectedly accumulated in skeletal muscle and liver. The basis of this divergence remains unclear but does not appear to correlate with ATG4A abundance, as ATG4A expression is not particularly elevated in these tissues (12) and likely reflects tissue-specific differences in autophagosome dynamics or mATG8 turnover. Mechanistically, impaired mATG8 priming, together with the absence of ATG4B/C/D, previously implicated in autophagosome formation (69), leads to the accumulation of unclosed pre-autophagosomal structures in ATG4A-only cells. As a result, fully formed autophagosomes (STX17TM-positive) are drastically reduced and autophagic flux is severely compromised, though not entirely abolished. The residual activity of ATG4A supports formation of a small number of functional autophagosomes, but this is insufficient to prevent the accumulation of p62/SQSTM1-positive aggregates, which appear both in cultured cells upon proteotoxic stress and in mouse tissues spontaneously. ATG4A-only cells also display reduced levels of ULK1, FIP200, ATG13 and ATG101, core components of the ULK1 initiation complex (46), consistent with the requirement of lipidated mATG8s for stabilizing this complex (73). These defects collectively place ATG4A-only cells in a state of chronic autophagy insufficiency that compromises proteostasis and stress resilience.

At the organismal level, ATG4A-only mice reveal the profound physiological consequences of severely restricted systemic autophagy, an experimental scenario that has been historically inaccessible because complete autophagy ablation is embryonic or perinatal lethal. Unlike heterozygous *Ambra1* or *Becn1* heterozygous mice, which show relatively mild autophagy reduction (74), ATG4A-only mice exhibit a deep impairment that manifests as early-onset multimorbidity: reduced body size, motor dysfunction, osteoporosis, tissue fibrosis, lymphopenia, and a systemic inflammatory state resembling inflammaging. At the molecular level, their hepatic transcriptome is defined by a coordinated upregulation of inflammatory and interferon-stimulated pathways together with a marked repression of somatotrophic and thyroid signaling. This scenario depicts a transcriptional rewiring that not only mirrors the inflammatory and metabolic disequilibrium typical of aged tissues but also aligns closely with molecular hallmarks consistently reported across independent aging datasets. In parallel, ATG4A-only mice exhibit elevated epigenetic age and multiple progeroid traits, including lipodystrophy, increased energy expenditure, GH–IGF-I axis dysregulation, hepatic glycogen accumulation, heightened DNA damage and robust activation of senescence markers. The strong enrichment of ATG4A-only differentially expressed genes among signatures of diverse progeroid models further reinforces the notion that systemic autophagy capacity is tightly coupled to the pace of aging in mammals. Notably, when considered alongside previously reported *Atg4*-deficient models, with *Atg4B* knockout mice showing only modest reductions in lifespan and *Atg4B/C* or *Atg4B/D* double mutants exhibiting intermediate survival defects, the dramatic shortening of ATG4A-only mice lifespan underscores a graded relationship between the depth of autophagy impairment and the severity of organismal decline. Together, these findings demonstrate that autophagic competence is a central determinant of organismal longevity and suggest that specific strategies aimed at preserving or enhancing autophagy may have broad therapeutic relevance for mitigating functional decline in both normal and premature aging.

## ACKNOWLEDGMENTS

This work was supported by grants from Ministerio de Ciencia e Innovación (Spain) (PID2021-127534OB-I00, PDI2020-118394RB-100 and PID2023-148089OB-I00) and Consejería de Ciencia, Innovación y Universidad del Gobierno del Principado de Asturias (AYUD/2021/51062 and IDE/2024/000784) and the South-Eastern Norway Regional Health Authority (2024072 to N.E.). The IUOPA is funded by the Asturian Government and Fundación Cajastur.

## AUTHOR CONTRIBUTIONS

G. G. M-G., M. F. S., I. T-G., Á.F.F., C.B., P.M., A.C-R., V.F., O.F.C., C.S., N. E. and G. M. performed experiments. Á.F.F., M.M.R-S. and G. M. participated in the generation, analysis, and maintenance of mice. A.A. performed histological analyses. P.M.Q., A.P.U., D.R-V., V.C-C. and J.M.P.F. performed RNA-Seq and epigenetic analyses. C. L-O. and G. M. devised the concept and supervised the project. G. M. wrote the manuscript.

## REFERENCES

1. Kuma A, Komatsu M, and Mizushima N. Autophagy-monitoring and autophagy-deficient mice. Autophagy. 2017;13(10):1619–28.

2. Marino G, Madeo F, and Kroemer G. Autophagy for tissue homeostasis and neuroprotection. Current Opinion in Cell Biology. 2011;23(2):198–206.

3. Rubinsztein DC, Marino G, and Kroemer G. Autophagy and aging. Cell. 2011;146(5):682–95.

4. Numan MS, Brown JP, and Michou L. Impact of air pollutants on oxidative stress in common autophagy-mediated aging diseases. International journal of environmental research and public health. 2015;12(2):2289–305.

5. Lopez-Otin C, Blasco MA, Partridge L, Serrano M, and Kroemer G. The hallmarks of aging. Cell. 2013;153(6):1194–217.

6. Lopez-Otin C, Blasco MA, Partridge L, Serrano M, and Kroemer G. Hallmarks of aging: An expanding universe. Cell. 2023;186(2):243–78.

7. Fernandez AF, and Lopez-Otin C. The functional and pathologic relevance of autophagy proteases. J Clin Invest. 2015;125(1):33–41.

8. Kirisako T, Ichimura Y, Okada H, Kabeya Y, Mizushima N, Yoshimori T, et al. The reversible modification regulates the membrane-binding state of Apg8/Aut7 essential for autophagy and the cytoplasm to vacuole targeting pathway. J Cell Biol. 2000;151(2):263–76.

9. Tanida I, Sou YS, Ezaki J, Minematsu-Ikeguchi N, Ueno T, and Kominami E. HsAtg4B/HsApg4B/autophagin-1 cleaves the carboxyl termini of three human Atg8 homologues and delipidates microtubule-associated protein light chain 3-and GABAA receptor-associated protein-phospholipid conjugates. J Biol Chem. 2004;279(35):36268–76.

10. Marino G, Uria JA, Puente XS, Quesada V, Bordallo J, and Lopez-Otin C. Human autophagins, a family of cysteine proteinases potentially implicated in cell degradation by autophagy. J Biol Chem. 2003;278(6):3671–8.

11. Tamargo-Gomez I, Martinez-Garcia GG, Suarez MF, Rey V, Fueyo A, Codina-Martinez H, et al. ATG4D is the main ATG8 delipidating enzyme in mammalian cells and protects against cerebellar neurodegeneration. Cell Death Differ. 2021;28(9):2651–72.

12. Tamargo-Gomez I, Martinez-Garcia GG, Suarez MF, Mayoral P, Bretones G, Astudillo A, et al. Analysis of ATG4C function in vivo. Autophagy. 2023;19(11):2912–33.

13. Cabrera S, Marino G, Fernandez AF, and Lopez-Otin C. Autophagy, proteases and the sense of balance. Autophagy. 2010;6(7):961–3.

14. Cabrera S, Fernandez AF, Marino G, Aguirre A, Suarez MF, Espanol Y, et al. ATG4B/autophagin-1 regulates intestinal homeostasis and protects mice from experimental colitis. Autophagy. 2013;9(8):1188–200.

15. Aguirre A, Lopez-Alonso I, Gonzalez-Lopez A, Amado-Rodriguez L, Batalla-Solis E, Astudillo A, et al. Defective autophagy impairs ATF3 activity and worsens lung injury during endotoxemia. J Mol Med (Berl). 2014;92(6):665–76.

16. Cabrera S, Maciel M, Herrera I, Nava T, Vergara F, Gaxiola M, et al. Essential role for the ATG4B protease and autophagy in bleomycin-induced pulmonary fibrosis. Autophagy. 2015;11(4):670–84.

17. Fernandez AF, Barcena C, Martinez-Garcia GG, Tamargo-Gomez I, Suarez MF, Pietrocola F, et al. Autophagy couteracts weight gain, lipotoxicity and pancreatic beta-cell death upon hypercaloric pro-diabetic regimens. Cell death & disease. 2017;8(8):e2970.

18. Martinez-Garcia GG, Perez RF, Fernandez AF, Durand S, Kroemer G, and Marino G. Autophagy Deficiency by Atg4B Loss Leads to Metabolomic Alterations in Mice. Metabolites. 2021;11(8).

19. Rodriguez-Muela N, Germain F, Marino G, Fitze PS, and Boya P. Autophagy promotes survival of retinal ganglion cells after optic nerve axotomy in mice. Cell Death Differ. 2012;19(1):162–9.

20. Marino G, Fernandez AF, Cabrera S, Lundberg YW, Cabanillas R, Rodriguez F, et al. Autophagy is essential for mouse sense of balance. The Journal of clinical investigation. 2010;120(7):2331–44.

21. Luhr M, Szalai P, and Engedal N. The Lactate Dehydrogenase Sequestration Assay - A Simple and Reliable Method to Determine Bulk Autophagic Sequestration Activity in Mammalian Cells. J Vis Exp. 2018(137).

22. Luhr M, Saetre F, and Engedal N. The Long-lived Protein Degradation Assay: an Efficient Method for Quantitative Determination of the Autophagic Flux of Endogenous Proteins in Adherent Cell Lines. Bio Protoc. 2018;8(9):e2836.

23. Parfitt AM, Drezner MK, Glorieux FH, Kanis JA, Malluche H, Meunier PJ, et al. Bone histomorphometry: standardization of nomenclature, symbols, and units. Report of the ASBMR Histomorphometry Nomenclature Committee. J Bone Miner Res. 1987;2(6):595–610.

24. Best L, Dost T, Esser D, Flor S, Haase M, Kadibalban AS, et al. Metabolic modeling reveals the aging-associated decline of host-microbiome metabolic interactions in mice. bioRxiv. 2024:2024.03.28.587009.

25. Benayoun BA, Pollina EA, Singh PP, Mahmoudi S, Harel I, Casey KM, et al. Remodeling of epigenome and transcriptome landscapes with aging in mice reveals widespread induction of inflammatory responses. Genome Res. 2019;29(4):697–709.

26. Zhou Q, Wan Q, Jiang Y, Liu J, Qiang L, and Sun L. A Landscape of Murine Long Non-Coding RNAs Reveals the Leading Transcriptome Alterations in Adipose Tissue during Aging. Cell Rep. 2020;31(8):107694.

27. Aramillo Irizar P, Schauble S, Esser D, Groth M, Frahm C, Priebe S, et al. Transcriptomic alterations during ageing reflect the shift from cancer to degenerative diseases in the elderly. Nat Commun. 2018;9(1):327.

28. Hahn O, Gronke S, Stubbs TM, Ficz G, Hendrich O, Krueger F, et al. Dietary restriction protects from age-associated DNA methylation and induces epigenetic reprogramming of lipid metabolism. Genome Biol. 2017;18(1):56.

29. Takasugi M, Nonaka Y, Takemura K, Yoshida Y, Stein F, Schwarz JJ, et al. An atlas of the aging mouse proteome reveals the features of age-related post-transcriptional dysregulation. Nat Commun. 2024;15(1):8520.

30. Stoeger T, Grant RA, McQuattie-Pimentel AC, Anekalla KR, Liu SS, Tejedor-Navarro H, et al. Aging is associated with a systemic length-associated transcriptome imbalance. Nat Aging. 2022;2(12):1191–206.

31. Schaum N, Lehallier B, Hahn O, Palovics R, Hosseinzadeh S, Lee SE, et al. Ageing hallmarks exhibit organ-specific temporal signatures. Nature. 2020;583(7817):596–602.

32. Schumacher B, van der Pluijm I, Moorhouse MJ, Kosteas T, Robinson AR, Suh Y, et al. Delayed and accelerated aging share common longevity assurance mechanisms. PLoS Genet. 2008;4(8):e1000161.

33. Osorio FG, Navarro CL, Cadinanos J, Lopez-Mejia IC, Quiros PM, Bartoli C, et al. Splicing-directed therapy in a new mouse model of human accelerated aging. Sci Transl Med. 2011;3(106):106ra7.

34. van der Pluijm I, Garinis GA, Brandt RM, Gorgels TG, Wijnhoven SW, Diderich KE, et al. Impaired genome maintenance suppresses the growth hormone--insulin-like growth factor 1 axis in mice with Cockayne syndrome. PLoS Biol. 2007;5(1):e2.

35. Ortega-Molina A, Lebrero-Fernandez C, Sanz A, Calvo-Rubio M, Deleyto-Seldas N, de Prado-Rivas L, et al. A mild increase in nutrient signaling to mTORC1 in mice leads to parenchymal damage, myeloid inflammation and shortened lifespan. Nat Aging. 2024;4(8):1102–20.

36. Lasry A, and Ben-Neriah Y. Senescence-associated inflammatory responses: aging and cancer perspectives. Trends Immunol. 2015;36(4):217–28.

37. De Cecco M, Ito T, Petrashen AP, Elias AE, Skvir NJ, Criscione SW, et al. L1 drives IFN in senescent cells and promotes age-associated inflammation. Nature. 2019;566(7742):73–8.

38. Barrett JE, Herzog CM, Aminzadeh-Gohari S, Redl E, Ishaq Parveen I, Rotharmel J, et al. Epigenetic signatures in surrogate tissues are able to assess cancer risk and indicate the efficacy of preventive measures. Commun Med (Lond). 2025;5(1):97.

39. Aryee MJ, Jaffe AE, Corrada-Bravo H, Ladd-Acosta C, Feinberg AP, Hansen KD, et al. Minfi: a flexible and comprehensive Bioconductor package for the analysis of Infinium DNA methylation microarrays. Bioinformatics. 2014;30(10):1363–9.

40. Pidsley R, CC YW, Volta M, Lunnon K, Mill J, and Schalkwyk LC. A data-driven approach to preprocessing Illumina 450K methylation array data. BMC Genomics. 2013;14:293.

41. Zipple MN, Zhao I, Kuo DC, Lee SM, Sheehan MJ, and Zhou W. Ecological Realism Accelerates Epigenetic Aging in Mice. Aging Cell. 2025;24(6):e70098.

42. Moqri M, Herzog C, Poganik JR, Biomarkers of Aging C, Justice J, Belsky DW, et al. Biomarkers of aging for the identification and evaluation of longevity interventions. Cell. 2023;186(18):3758–75.

43. Shvets E, Fass E, and Elazar Z. Utilizing flow cytometry to monitor autophagy in living mammalian cells. Autophagy. 2008;4(5):621–8.

44. Abreu S, Kriegenburg F, Gomez-Sanchez R, Mari M, Sanchez-Wandelmer J, Skytte Rasmussen M, et al. Conserved Atg8 recognition sites mediate Atg4 association with autophagosomal membranes and Atg8 deconjugation. EMBO Rep. 2017;18(5):765–80.

45. Itakura E, and Mizushima N. Characterization of autophagosome formation site by a hierarchical analysis of mammalian Atg proteins. Autophagy. 2010;6(6):764–76.

46. Yamamoto H, Zhang S, and Mizushima N. Autophagy genes in biology and disease. Nat Rev Genet. 2023;24(6):382–400.

47. Koyama-Honda I, Itakura E, Fujiwara TK, and Mizushima N. Temporal analysis of recruitment of mammalian ATG proteins to the autophagosome formation site. Autophagy. 2013;9(10):1491–9.

48. Marino G, Fernandez AF, Cabrera S, Lundberg YW, Cabanillas R, Rodriguez F, et al. Autophagy is essential for mouse sense of balance. Journal of Clinical Investigation. 2010;120(7):2331–44.

49. Itakura E, Kishi-Itakura C, and Mizushima N. The hairpin-type tail-anchored SNARE syntaxin 17 targets to autophagosomes for fusion with endosomes/lysosomes. Cell. 2012;151(6):1256–69.

50. Ezaki J, Matsumoto N, Takeda-Ezaki M, Komatsu M, Takahashi K, Hiraoka Y, et al. Liver autophagy contributes to the maintenance of blood glucose and amino acid levels. Autophagy. 2011;7(7):727–36.

51. Harkema L, Youssef SA, and de Bruin A. Pathology of Mouse Models of Accelerated Aging. Vet Pathol. 2016;53(2):366–89.

52. Marino G, Ugalde AP, Fernandez AF, Osorio FG, Fueyo A, Freije JMP, et al. Insulin-like growth factor 1 treatment extends longevity in a mouse model of human premature aging by restoring somatotroph axis function. Proceedings of the National Academy of Sciences of the United States of America. 2010;107(37):16268–73.

53. Garinis GA, van der Horst GT, Vijg J, and Hoeijmakers JH. DNA damage and ageing: new-age ideas for an age-old problem. Nat Cell Biol. 2008;10(11):1241–7.

54. Marino G, Ugalde AP, Salvador-Montoliu N, Varela I, Quiros PM, Cadinanos J, et al. Premature aging in mice activates a systemic metabolic response involving autophagy induction. Human Molecular Genetics. 2008;17(14):2196–211.

55. Niedernhofer LJ, Garinis GA, Raams A, Lalai AS, Robinson AR, Appeldoorn E, et al. A new progeroid syndrome reveals that genotoxic stress suppresses the somatotroph axis. Nature. 2006;444(7122):1038–43.

56. Barcena C, Quiros PM, Durand S, Mayoral P, Rodriguez F, Caravia XM, et al. Methionine Restriction Extends Lifespan in Progeroid Mice and Alters Lipid and Bile Acid Metabolism. Cell Rep. 2018;24(9):2392–403.

57. Lopez-Mejia IC, de Toledo M, Chavey C, Lapasset L, Cavelier P, Lopez-Herrera C, et al. Antagonistic functions of LMNA isoforms in energy expenditure and lifespan. EMBO Rep. 2014;15(5):529–39.

58. Schefer V, and Talan MI. Oxygen consumption in adult and AGED C57BL/6J mice during acute treadmill exercise of different intensity. Exp Gerontol. 1996;31(3):387–92.

59. Penn DJ, Zala SM, and Luzynski KC. Regulation of Sexually Dimorphic Expression of Major Urinary Proteins. Front Physiol. 2022;13:822073.

60. Garratt M, Stockley P, Armstrong SD, Beynon RJ, and Hurst JL. The scent of senescence: sexual signalling and female preference in house mice. J Evol Biol. 2011;24(11):2398–409.

61. Yan J, Gao Y, Bai J, Li J, Li M, Liu X, et al. SERPINB1 promotes Senecavirus A replication by degrading IKBKE and regulating the IFN pathway via autophagy. J Virol. 2023;97(10):e0104523.

62. Barnhoorn S, Meima ME, Peeters RP, Darras VM, Leeuwenburgh S, Hoeijmakers JHJ, et al. Decreased hepatic thyroid hormone signaling in systemic and liver-specific but not brain-specific accelerated aging due to DNA repair deficiency in mice. Eur Thyroid J. 2023;12(6).

63. Visser WE, Bombardieri CR, Zevenbergen C, Barnhoorn S, Ottaviani A, van der Pluijm I, et al. Tissue-Specific Suppression of Thyroid Hormone Signaling in Various Mouse Models of Aging. PLoS One. 2016;11(3):e0149941.

64. Wiechert L, Nemeth J, Pusterla T, Bauer C, De Ponti A, Manthey S, et al. Hepatocyte-specific S100a8 and S100a9 transgene expression in mice causes Cxcl1 induction and systemic neutrophil enrichment. Cell Commun Signal. 2012;10(1):40.

65. Gupta U, Ghosh S, Wallace CT, Shang P, Xin Y, Nair AP, et al. Increased LCN2 (lipocalin 2) in the RPE decreases autophagy and activates inflammasome-ferroptosis processes in a mouse model of dry AMD. Autophagy. 2023;19(1):92–111.

66. Asimakopoulou A, Weiskirchen S, and Weiskirchen R. Lipocalin 2 (LCN2) Expression in Hepatic Malfunction and Therapy. Front Physiol. 2016;7:430.

67. Carrero D, Soria-Valles C, and Lopez-Otin C. Hallmarks of progeroid syndromes: lessons from mice and reprogrammed cells. Dis Model Mech. 2016;9(7):719–35.

68. Collado M, Blasco MA, and Serrano M. Cellular senescence in cancer and aging. Cell. 2007;130(2):223–33.

69. Kauffman KJ, Yu S, Jin J, Mugo B, Nguyen N, O’Brien A, et al. Delipidation of mammalian Atg8-family proteins by each of the four ATG4 proteases. Autophagy. 2018;14(6):992–1010.

70. Nguyen TN, Padman BS, Zellner S, Khuu G, Uoselis L, Lam WK, et al. ATG4 family proteins drive phagophore growth independently of the LC3/GABARAP lipidation system. Mol Cell. 2021;81(9):2013–30 e9.

71. Agrotis A, Pengo N, Burden JJ, and Ketteler R. Redundancy of human ATG4 protease isoforms in autophagy and LC3/GABARAP processing revealed in cells. Autophagy. 2019;15(6):976–97.

72. Nguyen N, Olivas TJ, Mires A, Jin J, Yu S, Luan L, et al. The insufficiency of ATG4A in macroautophagy. J Biol Chem. 2020;295(39):13584–600.

73. Grunwald DS, Otto NM, Park JM, Song D, and Kim DH. GABARAPs and LC3s have opposite roles in regulating ULK1 for autophagy induction. Autophagy. 2020;16(4):600–14.

74. Ramirez-Pardo I, Villarejo-Zori B, Jimenez-Loygorri JI, Sierra-Filardi E, Alonso-Gil S, Marino G, et al. Ambra1 haploinsufficiency in CD1 mice results in metabolic alterations and exacerbates age-associated retinal degeneration. Autophagy. 2023;19(3):784–804.

