## Supplementary Information for "Combined loss of ATG4B, ATG4C and ATG4D impairs systemic autophagy and triggers accelerated aging in mice"

### SUPPLEMENTARY FIGURE LEGENDS

**Figure S1: Alterations in the relative size of organs from ATG4A-only mice. (A)** Representative images and relative size (wet weight/body weight) quantification of different organs from WT and ATG4A-only mice.

**Figure S2. Analysis of oxygen production and carbon dioxide production in WT and ATG4A-only mice.** Line plots represent mean ± SEM. *, $P < 0.05$; Wilcoxon signed-rank test (24 h), linear models with cluster-robust standard errors (day/night). (N = 5 for and 6 for ATG4A-only mice).

**Figure S3. PCA plot of the analysis of ATG4A-only versus WT liver mouse transcriptomes.** Each sample is represented on the plot based on principal components 1 and 2. Color groups indicate the sex and genotype of each sample.

**Figure S4. Heatmap of leading-edge genes of selected gene sets.** Each panel depicts columns corresponding to the different samples analyzed (grouped by sex and genotype), and rows representing the leading-edge genes identified in the respective GSEA analysis. Genes are clustered before representation. Cell colors indicate row-scaled gene expression levels.

**Figure S5. Heatmap of leading-edge genes of WT-enriched gene sets.** Each panel depicts columns corresponding to the different samples analyzed (grouped by sex and genotype), and rows representing the leading-edge genes identified in the respective GSEA analysis. Genes are clustered before representation. Cell colors indicate row-scaled gene expression levels.

**Figure S6. Running score plots of Hallmarks MSigDB gene sets. A)** ATG4A-only-enriched gene sets when compared to age-matched WT mice transcriptomes. **B)** ATG4A-only-depleted gene sets when compared to age-matched WT mice transcriptomes.

**Figure S7. Running score plots of progeria models gene sets. A)** ATG4A-only-enriched gene sets that are either convergently enriched or convergently depleted (last gene set) in different progeria models when compared to age-matched WT mice transcriptomes. **B)** ATG4A-only-enriched senescence gene sets when compared to age-matched WT mice transcriptomes

**Figure S8. GABARAP and GABARAPL1 lipidation in ATG4A-only and ATG4 single and double knockout MEFs.** Representative immunoblotting of GABARAP and GABARAPL1 proteins in protein extracts from MEFs of the indicated genotypes cultured in DMEM (control conditions). LOAD: β-actin.

**Table S1. RNA-Seq analysis of gene expression in 4 month-old WT and ATG4A-only mouse livers.** Total detected transcripts for which different parameters in each mouse liver sample is shown, namely, **baseMean**: average of DESeq2-normalized count variables for a specific gene across all samples; **$log_2$FoldChange**: DESeq2-calculated $log_2$ Fold Change in gene expression between TKO and WT mice; lfcSE: standard error estimate for the $log_2$ Fold Change; stat: difference in deviance between the reduced model and the full model; **pvalue**: DESeq2-calculated p value for the likelihood ratio test (LRT) of full vs. reduced model; **padj**: Benjamini-Hochberg adjusted p-value.

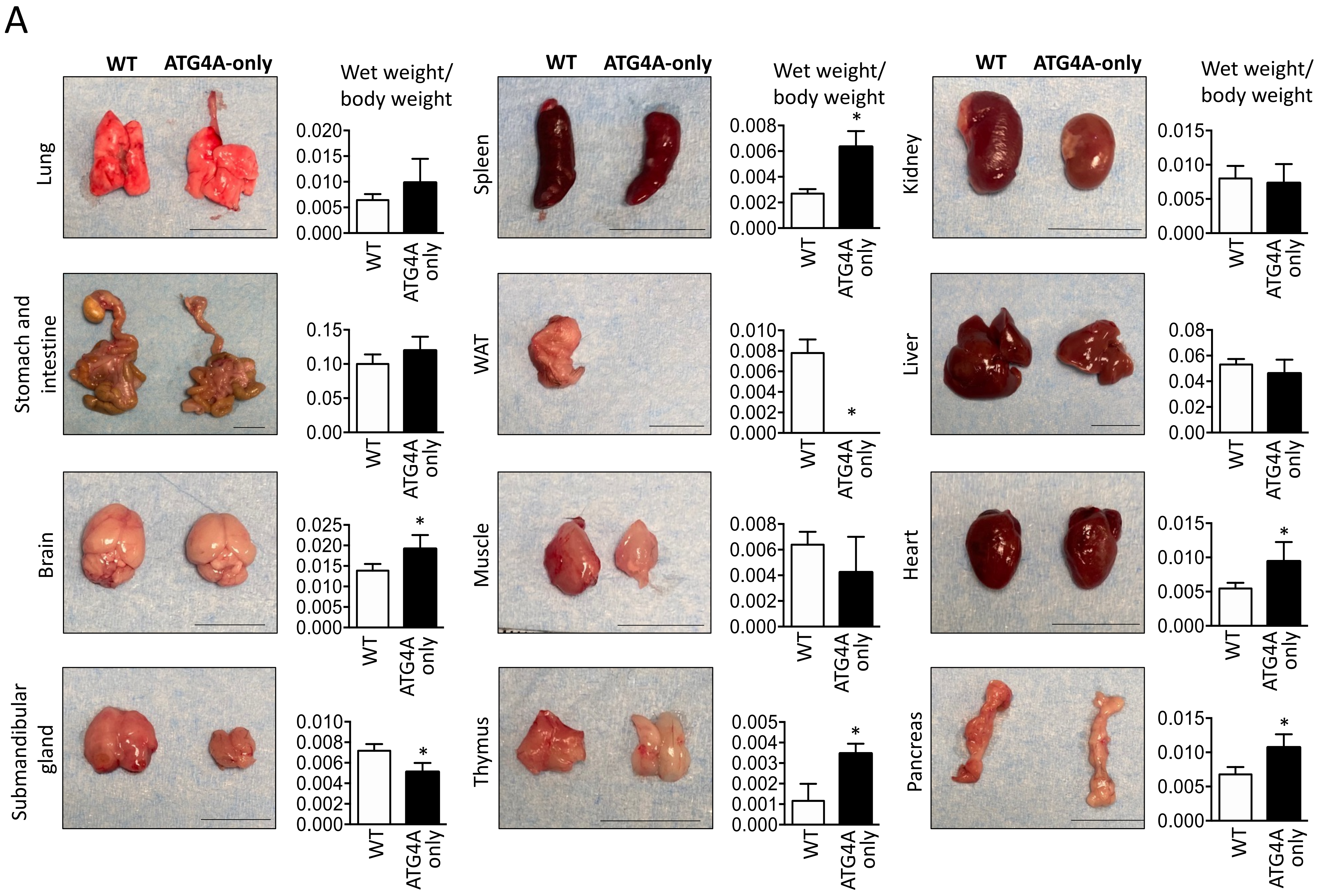

A
WT
ATG4A-only
Wet weight/ body weight
Lung
Spleen
Kidney
Stomach and intestine
WAT
Liver
Brain
Muscle
Heart
Submandibular gland
Thymus
Pancreas
WT
ATG4A only
*


Figure S1

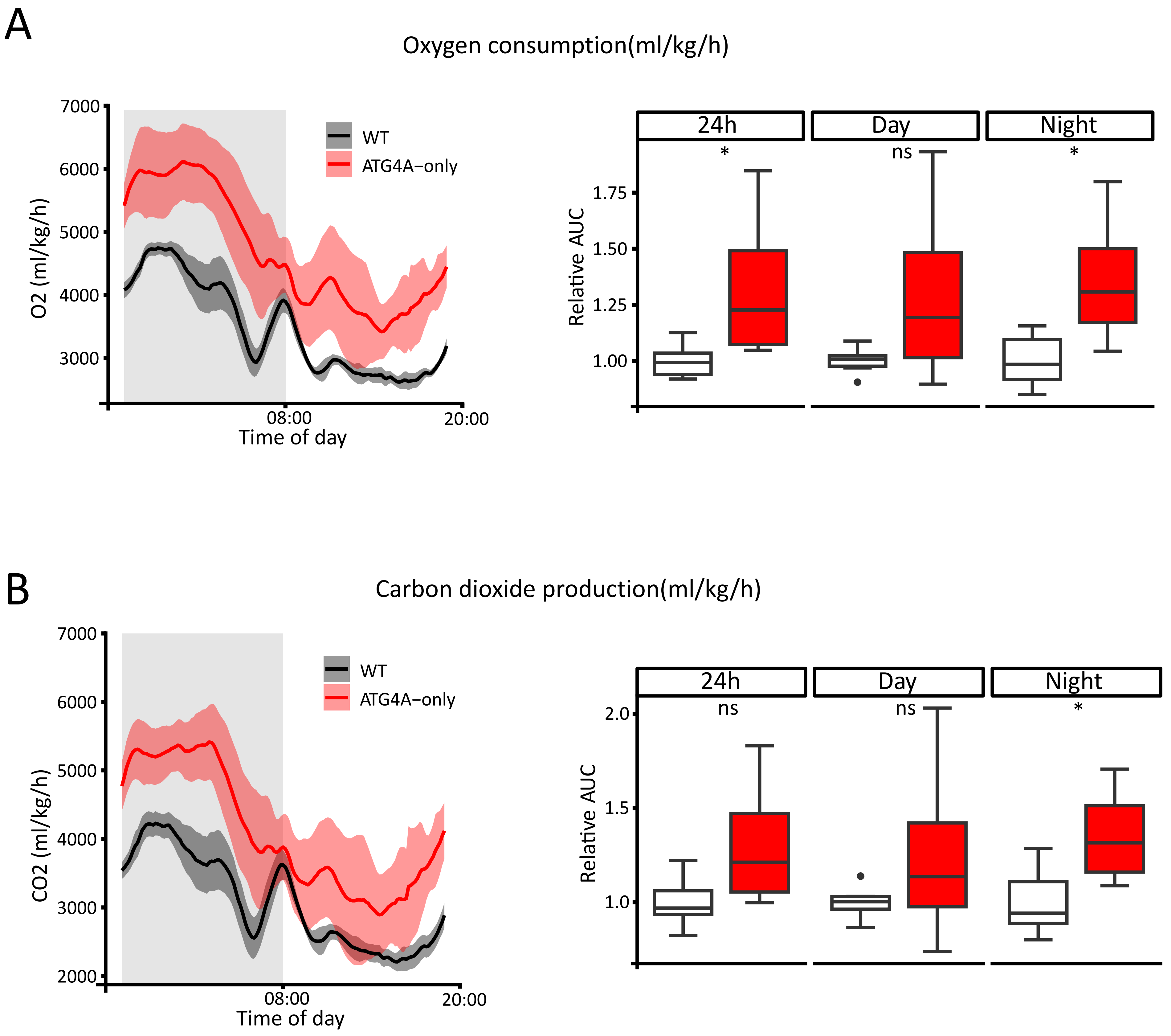
A
Oxygen consumption(ml/kg/h)
O2 (ml/kg/h)
7000
6000
5000
4000
3000
08:00
20:00
Time of day
WT
ATG4A−only
24h
Day
Night
*
ns
*
Relative AUC
1.75
1.50
1.25
1.00
B
Carbon dioxide production(ml/kg/h)
CO2 (ml/kg/h)
7000
6000
5000
4000
3000
2000
08:00
20:00
Time of day
WT
ATG4A−only
24h
Day
Night
ns
ns
*
Relative AUC
2.0
1.5
1.0

Figure S2

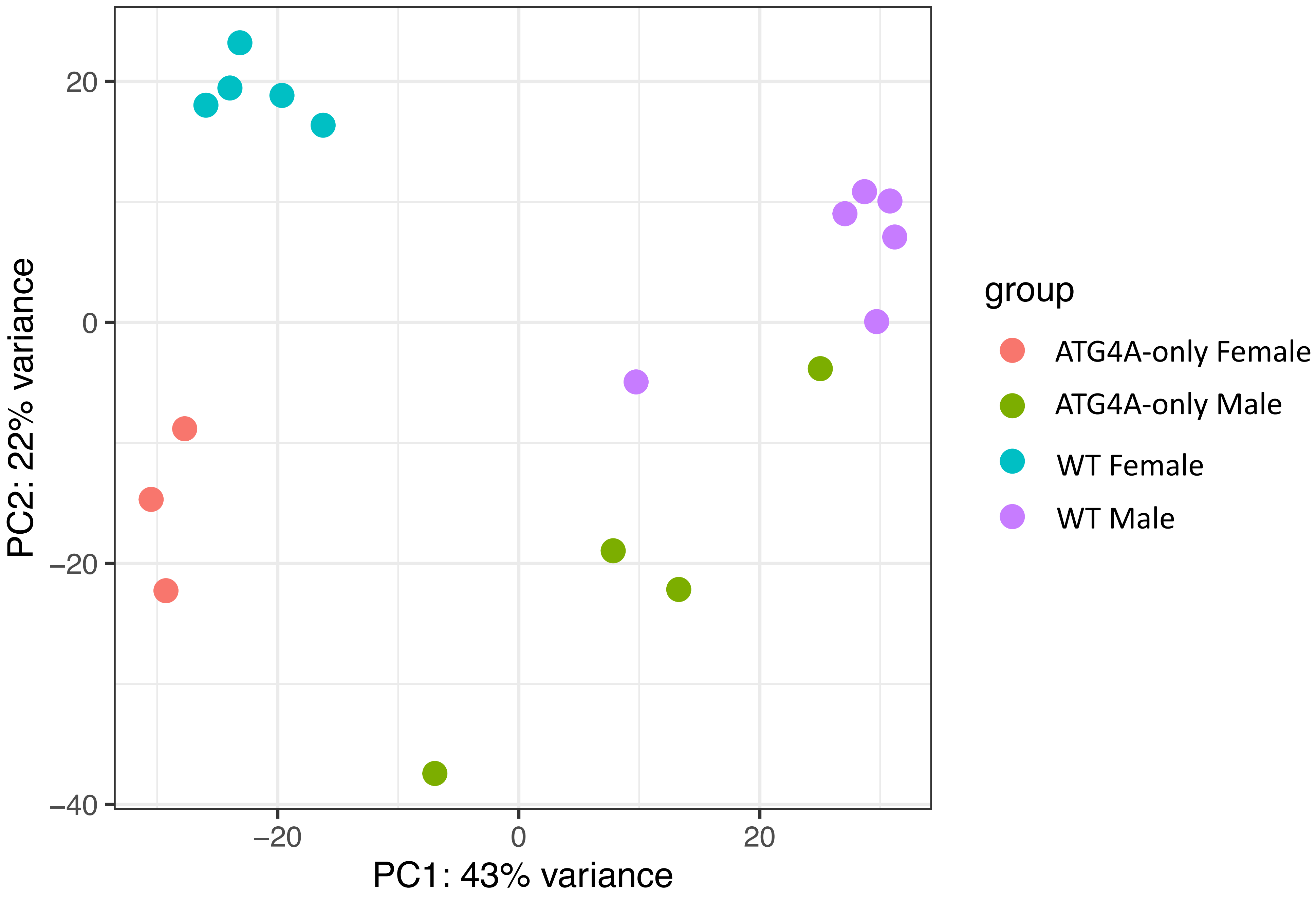
PC2: 22% variance
PC1: 43% variance
20
0
−20
−40
−20
0
20
group
ATG4A-only Female
ATG4A-only Male
WT Female
WT Male

Figure S3

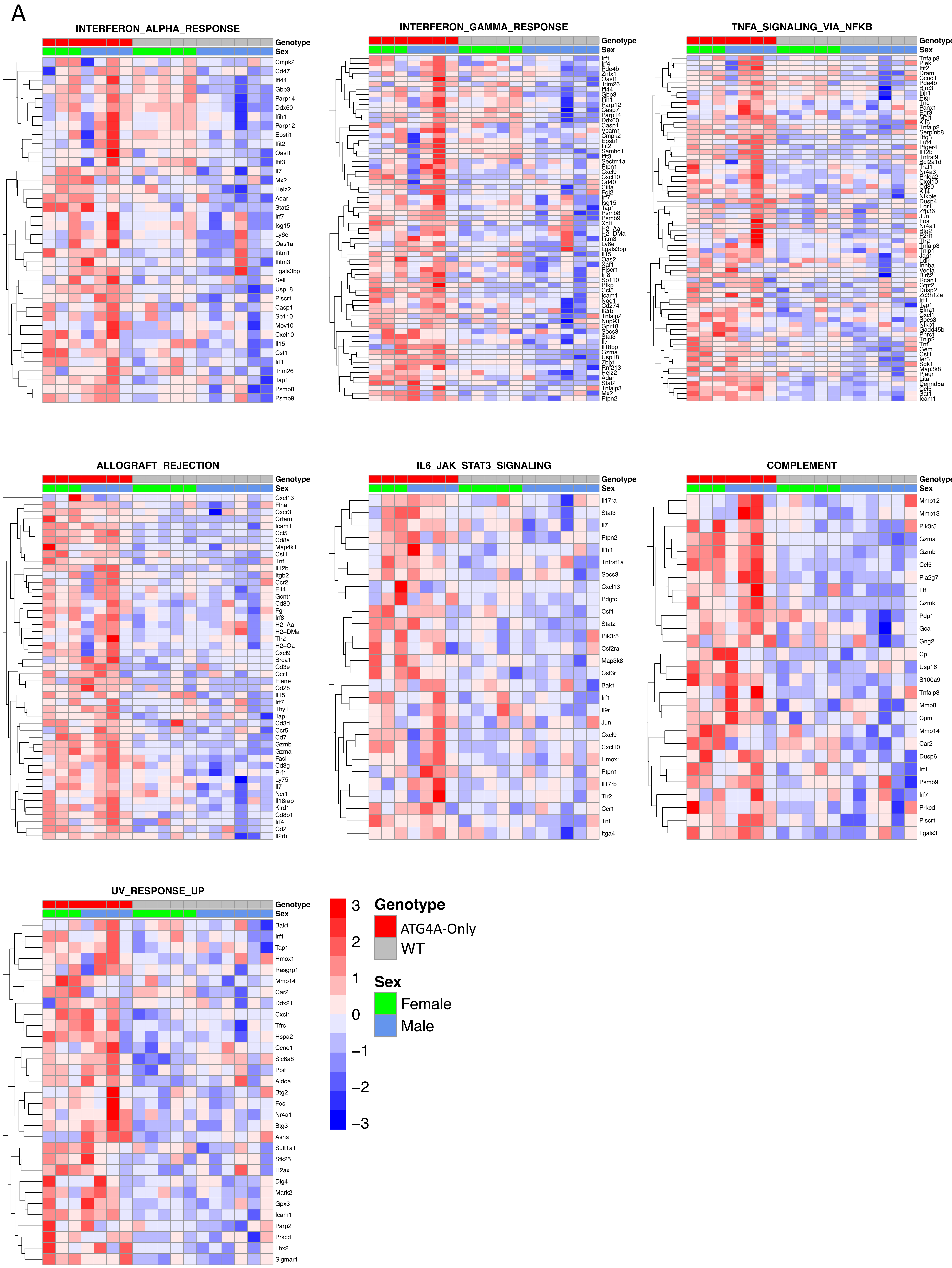
A
INTERFERON_ALPHA_RESPONSE
Genotype
Sex
Cmpk2
Cd47
Ifi44
Gbp3
Parp14
Ddx60
Ifih1
Parp12
Epsti1
Ifit2
Oasl1
Ifit3
Il7
Mx2
Helz2
Adar
Stat2
Irf7
Isg15
Ly6e
Oas1a
Ifitm1
Ifitm3
Lgals3bp
Sell
Usp18
Plscr1
Casp1
Sp110
Mov10
Cxcl10
Il15
Csf1
Irf1
Trim26
Tap1
Psmb8
Psmb9
INTERFERON_GAMMA_RESPONSE
TNFA_SIGNALING_VIA_NFKB
ALLOGRAFT_REJECTION
Cxcl13
Flna
Cxcr3
Crtam
Icam1
Ccl5
Cd8a
Map4k1
Csf1
Tnf
Il12b
Itgb2
Ccr2
Elf4
Gcnt1
Cd80
Fgr
Irf8
H2-Aa
H2-DMa
Tlr2
H2-Oa
Cxcl9
Brca1
Cd3e
Ccr1
Elane
Cd28
Il15
Irf7
Thy1
Tap1
Cd3d
Ccr5
Cd7
Gzmb
Gzma
Fasl
Cd3g
Prf1
Ly75
Il7
Ncr1
Il18rap
Klrd1
Cd8b1
Irf4
Cd2
Il2rb
IL6_JAK_STAT3_SIGNALING
Il17ra
Stat3
Il7
Ptpn2
Il1r1
Tnfrsf1a
Socs3
Cxcl13
Pdgfc
Csf1
Stat2
Pik3r5
Csf2ra
Map3k8
Csf3r
Bak1
Irf1
Il9r
Jun
Cxcl9
Cxcl10
Hmox1
Ptpn1
Il17rb
Tlr2
Ccr1
Tnf
Itga4
COMPLEMENT
Mmp12
Mmp13
Pik3r5
Gzma
Gzmb
Ccl5
Pla2g7
Ltf
Gzmk
Pdp1
Gca
Gng2
Cp
Usp16
S100a9
Tnfaip3
Mmp8
Cpm
Mmp14
Car2
Dusp6
Irf1
Psmb9
Irf7
Prkcd
Plscr1
Lgals3
UV_RESPONSE_UP
Bak1
Irf1
Tap1
Hmox1
Rasgrp1
Mmp14
Car2
Ddx21
Cxcl1
Tfrc
Hspa2
Ccne1
Slc6a8
Ppif
Aldoa
Btg2
Fos
Nr4a1
Btg3
Asns
Sult1a1
Stk25
H2ax
Dlg4
Mark2
Gpx3
Icam1
Parp2
Prkcd
Lhx2
Sigmar1
3
2
1
0
−1
−2
−3
Genotype
ATG4A-Only
WT
Sex
Female
Male

Figure S4

A

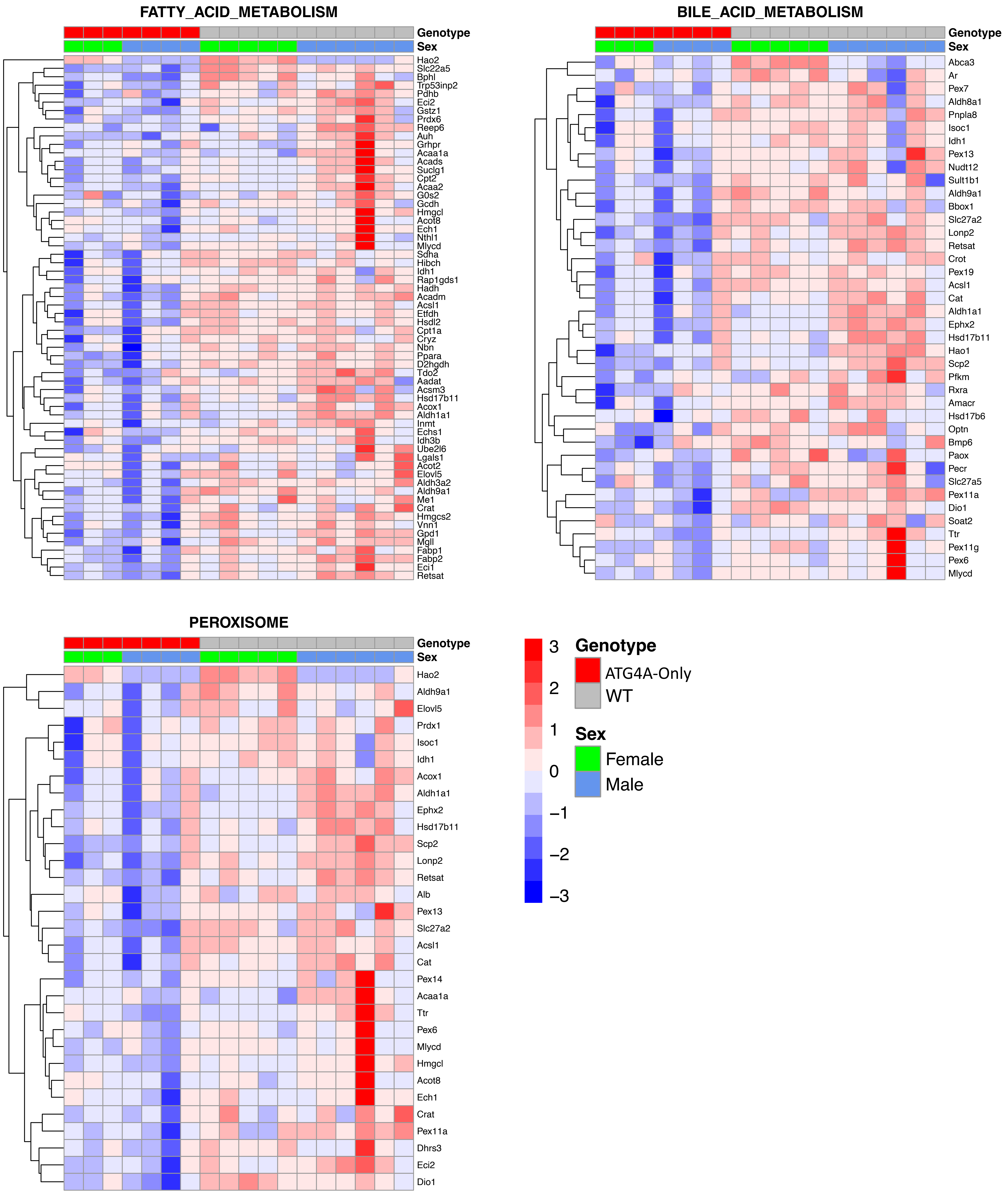
FATTY_ACID_METABOLISM
Genotype
Sex
Hao2
Slc22a5
Bphl
Trp53inp2
Pdhb
Eci2
Gstz1
Prdx6
Reep6
Auh
Grhpr
Acaa1a
Acads
Suclg1
Cpt2
Acaa2
G0s2
Gcdh
Hmgcl
Acot8
Ech1
Nthl1
Mlycd
Sdha
Hibch
Idh1
Rap1gds1
Hadh
Acadm
Acsl1
Etfdh
Hsdl2
Cpt1a
Cryz
Nbn
Ppara
D2hgdh
Tdo2
Aadat
Acsm3
Hsd17b11
Acox1
Aldh1a1
Inmt
Echs1
Idh3b
Ube2l6
Lgals1
Acot2
Elovl5
Aldh3a2
Aldh9a1
Me1
Crat
Hmgcs2
Vnn1
Gpd1
Mgll
Fabp1
Fabp2
Eci1
Retsat
BILE_ACID_METABOLISM
Genotype
Sex
Abca3
Ar
Pex7
Aldh8a1
Pnpla8
Isoc1
Idh1
Pex13
Nudt12
Sult1b1
Aldh9a1
Bbox1
Slc27a2
Lonp2
Retsat
Crot
Pex19
Acsl1
Cat
Aldh1a1
Ephx2
Hsd17b11
Hao1
Scp2
Pfkm
Rxra
Amacr
Hsd17b6
Optn
Bmp6
Paox
Pecr
Slc27a5
Pex11a
Dio1
Soat2
Ttr
Pex11g
Pex6
Mlycd
PEROXISOME
Genotype
Sex
Hao2
Aldh9a1
Elovl5
Prdx1
Isoc1
Idh1
Acox1
Aldh1a1
Ephx2
Hsd17b11
Scp2
Lonp2
Retsat
Alb
Pex13
Slc27a2
Acsl1
Cat
Pex14
Acaa1a
Ttr
Pex6
Mlycd
Hmgcl
Acot8
Ech1
Crat
Pex11a
Dhrs3
Eci2
Dio1
3
2
1
0
−1
−2
−3
Genotype
ATG4A-Only
WT
Sex
Female
Male

Figure S5

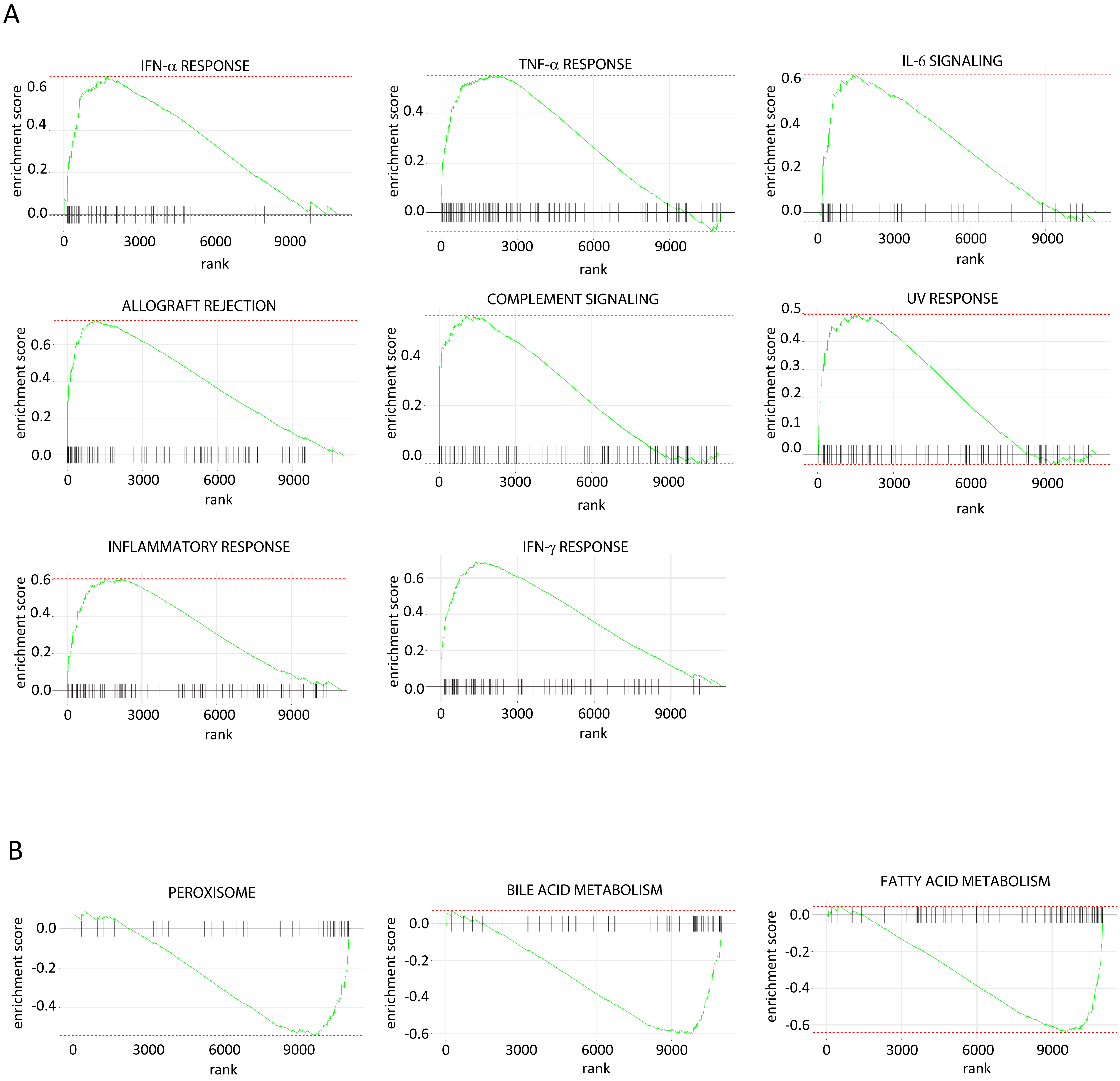
A
IFN-α RESPONSE
TNF-α RESPONSE
IL-6 SIGNALING
ALLOGRAFT REJECTION
COMPLEMENT SIGNALING
UV RESPONSE
INFLAMMATORY RESPONSE
IFN-γ RESPONSE
B
PEROXISOME
BILE ACID METABOLISM
FATTY ACID METABOLISM
enrichment score
rank
0
3000
6000
9000

Figure S6

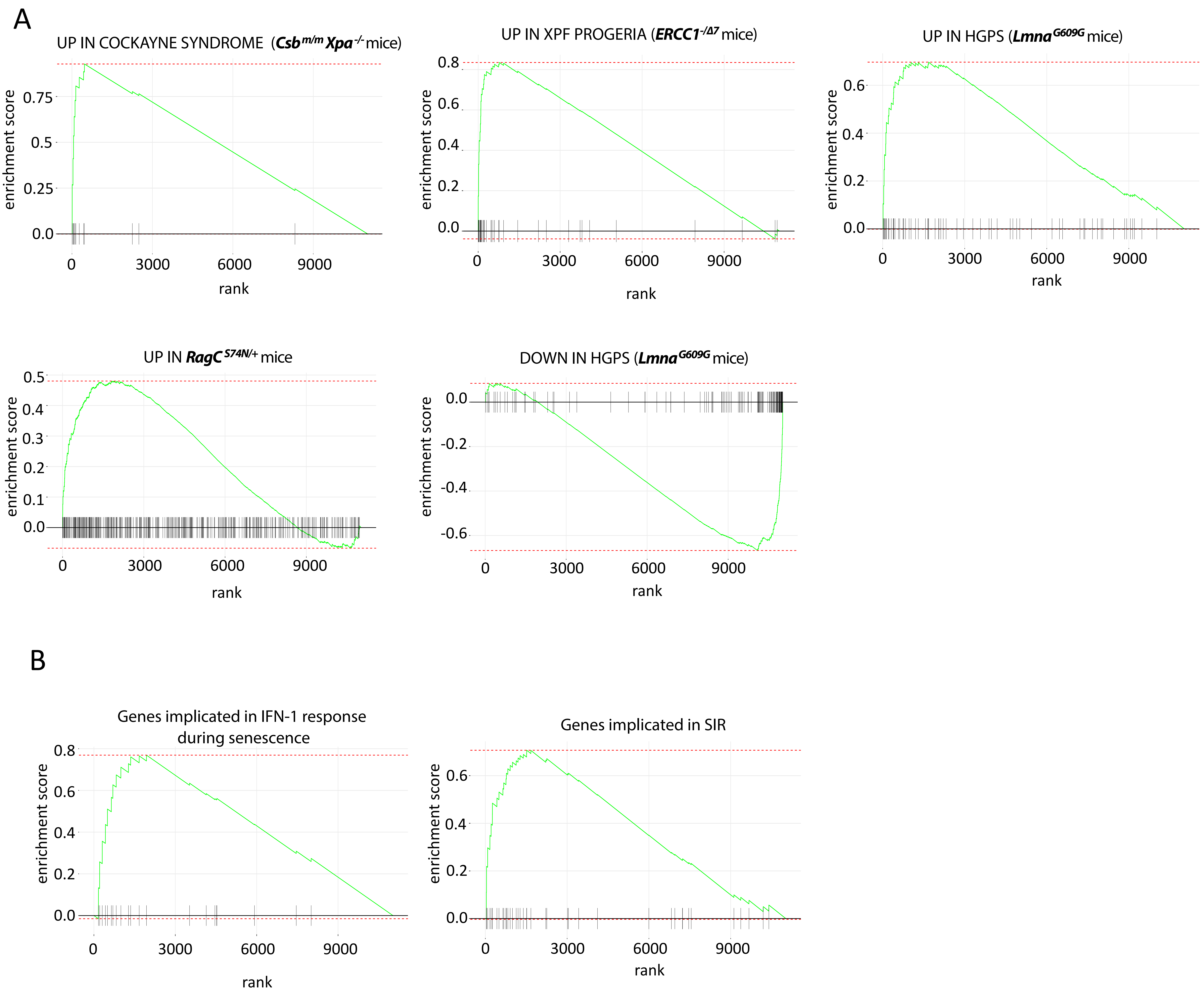
A
UP IN COCKAYNE SYNDROME (Csb$^{m/m}$ Xpa$^{-/-}$ mice)
UP IN XPF PROGERIA (ERCC1$^{-/\Delta 7}$ mice)
UP IN HGPS (Lmna$^{G609G}$ mice)
UP IN RagC$^{S74N/+}$ mice
DOWN IN HGPS (Lmna$^{G609G}$ mice)
enrichment score
rank
B
Genes implicated in IFN-1 response during senescence
Genes implicated in SIR
enrichment score
rank

Figure S7

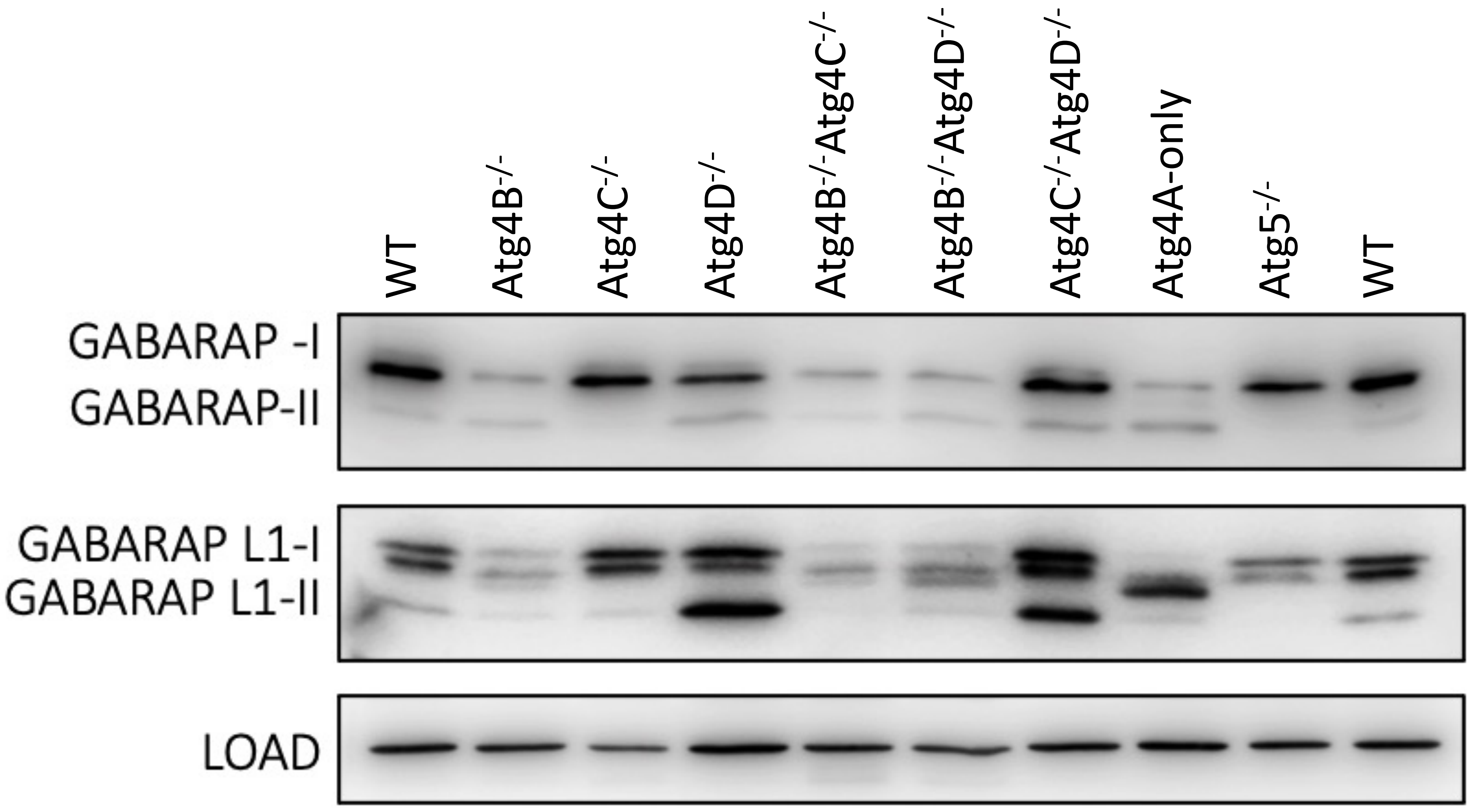
WT
Atg4B-/-
Atg4C-/-
Atg4D-/-
Atg4B-/-Atg4C-/-
Atg4B-/-Atg4D-/-
Atg4C-/-Atg4D-/-
Atg4A-only
Atg5-/-
WT
GABARAP -I
GABARAP-II
GABARAP L1-I
GABARAP L1-II
LOAD

Figure S8
